# The actin nucleator Spir-1 is a virus restriction factor that promotes IRF3 activation

**DOI:** 10.1101/2020.08.31.276659

**Authors:** Alice Abreu Torres, Stephanie L. Macilwee, Amir Rashid, Sarah E. Cox, Jonas D. Albarnaz, Claudio A. Bonjardim, Geoffrey L. Smith

## Abstract

Cellular proteins often have multiple and diverse functions. This is illustrated with protein Spir-1 that is an actin nucleator, but, as shown here, also functions to enhance IRF3 activation downstream of RNA sensing by RIG-I/MDA-5. In human and mouse cells lacking Spir-1, IRF3 activation is impaired, whereas Spir-1 overexpression enhanced IRF3 activation. Furthermore in Spir-1^-/-^ cells, the infectious virus titres and sizes of plaques formed by two viruses that are sensed by RIG-I, vaccinia virus (VACV) and Zika virus, are increased. These observations demonstrate the biological importance of Spir-1 in the response to virus infection. Like cellular proteins, viral proteins also have multiple and diverse functions. Here, we also show that VACV virulence factor K7 binds directly to Spir-1 and that a diphenylalanine motif of Spir-1 is needed for this interaction and for Spir-1-mediated enhancement of IRF3 activation. Thus, Spir-1 is a new virus restriction factor and is targeted directly by an immunomodulatory viral protein that enhances virus virulence and diminishes IRF3 activation.

**Author Summary:** Infection of cells by viruses is sensed by host molecules called pattern recognition receptors (PRRs) that activate signalling pathways leading to an anti-viral response. In turn, viruses express proteins that negate these host responses to mediate escape from the anti-viral response. Here, we report that protein K7 from a large DNA virus called vaccinia virus (VACV), binds to a host cell protein called Spir-1. Spir-1 is known to regulate the assembly of actin filaments inside cells, but here we show that Spir-1 also functions to activate the host response to virus infection and to limit the replication and spread of both RNA and DNA viruses. Thus, this study has uncovered new functions of cellular protein Spir-1 as an activator of innate immunity and as a restriction factor for diverse viruses. Further, it shows that Spir-1 is targeted by a virus protein during infection.

## Introduction

The host innate immune response to viral infection begins with the sensing of pathogen-associated molecular patterns (PAMPs) by pattern recognition receptors (PRRs), such as Toll-like receptors and retinoic acid-inducible gene-I (RIG-I)-like receptors (RLRs) (1, 2). The sensing of virus macromolecules, such as viral DNA or RNA, by PRRs triggers signalling cascades that culminate in the activation of the transcriptional factors interferon regulatory factor (IRF) 3, activator protein (AP) 1 and nuclear factor kappa B (NF-κB) (3). These transcriptional factors translocate to the nucleus where they induce the transcription of genes encoding interferons (IFN), cytokines and chemokines. Once secreted from the cell, IFNs, cytokines and chemokines promote inflammation to restrict virus replication and control the infection. IFNs bind to their receptors on the surface of infected or non-infected cells, and trigger signal transduction via the JAK-STAT pathway leading to expression of interferon-stimulated genes (ISGs) that induce an antiviral state (4).

During the co-evolution with their hosts, viruses have evolved strategies to evade, suppress or exploit the host response to infection by targeting multiple steps of the host immune response (5). Vaccinia virus (VACV), the prototypical orthopoxvirus, is well known as the live vaccine used to eradicate smallpox (6). VACV has a large dsDNA genome of approximately 190 kbp (7), replicates in the cytoplasm (8) and encodes scores of proteins that antagonise innate immunity (9). Interestingly, some cellular pathways, such as those leading to activation of IRF3 or NF-κB, or the JAK-STAT pathway downstream of IFNs binding to their receptors, are targeted by multiple different VACV proteins. Moreover, some of these viral antagonists are multifunctional and inhibit more than one host innate immune pathway (9).

VACV protein K7 is one such antagonist of innate immunity. K7 is a small, intracellular protein that is non-essential for virus replication in cell culture, yet contributes to virulence in both intradermal and intranasal mouse models of infection (10). Functionally, K7 was reported to suppress NF-κB activation by binding to interleukin-1 receptor-associated kinase-like 2 (IRAK2) and tumour necrosis factor (TNF) receptor-associated factor 6 (TRAF6) (11). K7 also inhibits IRF3 activation and binds to the DEAD-box RNA helicase 3 (DDX3) (11). Another study reported that K7 affected regulation of histone methylation during VACV infection, by an unknown mechanism (12). Two unbiased proteomic searches identified cellular binding partners of K7 (13, 14), suggesting that K7 may have additional functions. The identification of cellular proteins targeted by viral proteins has been a useful approach to identify cellular factors that function in the recognition and restriction of virus infections (15, 16). In this study, by investigating the interaction between K7 and the cellular protein Spir-1 (14), we identified new functions for Spir-1 as an activator of innate immunity and as a restriction factor for both DNA and RNA viruses.

The protein spire homolog 1 (Spir-1, also known as SPIRE1) was first described to affect *Drosophila* embryogenesis (17). Spir-1 has actin-binding domains (18) through which it nucleates actin filaments, an activity shared with the Arp2/3 complex and the formins (19). Spir-1 is organised in multiple functional domains. The N-terminal region contains the kinase non-catalytic C-lobe domain (KIND), which mediates Spir-1 interaction with other proteins, such as the formins (20). Spir-1 and the formins cooperate during actin nucleation (21-29). The KIND domain is followed by four actin-binding Wiskott-Andrich syndrome protein homology domain 2 (WH2) domains that are responsible for actin nucleation (19, 30, 31). The C-terminal region of Spir-1 contains a globular tail domain-binding motif (GTBM), which is responsible for binding to myosin V (32). Next, there is a Spir-box (SB) domain that is conserved within the Spir protein family. Due to its similarity to a helical region of the rabphilin-3A protein that interacts with the GTPase Rab3A, it is thought that the Spir-box domain is involved in the association of Spir-1 and Rab-GTPases (24, 33-35). Following the SB domain, there is a modified FYVE zinc finger domain that interacts with negatively-charged lipids in membranes. The membrane targeting specificity is mediated via the interaction of Spir-1 with other membrane-bound proteins such as the Rab-GTPases (29).

In adult mice, Spir-1 is expressed preferentially in neuronal and hematopoietic cells (36, 37) and in humans the brain also has high Spir-1 expression (38). In general, Spir-1 is involved in several actin-dependent cellular functions such as vesicle trafficking (24, 34, 35, 39), DNA repair (40), mitochondrial division (41), and development of germ cells (17, 23, 42) but a role in innate immunity has not been described. Interestingly, a genome-wide association study found a single nucleotide polymorphism in Spir-1 that correlated with a different antibody response to smallpox vaccination (43).

Here, Spir-1 is shown to promote IRF3 activation and this activity depends upon a diphenylalanine motif that is also necessary for the direct interaction of Spir-1 with VACV virulence factor K7. Using gain-of-function and loss-of-function cell lines, Spir-1 is shown to diminish VACV and ZIKV replication and/or spread and is therefore a viral restriction factor.

## Results

### Vaccinia virus protein K7 co-precipitates with the C terminal region of Spir-1

A previous proteomic study identified Spir-1 as a cellular interacting partner of VACV protein K7 although this interaction was not validated (14). Mammals have two *Spire* genes, *Spire1* and *Spire2* that encode closely related proteins (overall 42% identity and 58% similarity in humans), especially within the WH2 and Spir-box domains (37, 44). The Uniprot database for human Spir-1 (Q08AE8) describes five Spir-1 isoforms and splicing before the GTBM (Exon 9) and SB (Exon 13) domains has been demonstrated (45). Among Spir-1 isoforms, Spir-1 known as isoform 2 that doesn’t contain Exon 9 or Exon 13, is the most abundant form in the brain and small intestine tissues (45) and is the form studied here. To confirm if K7 co-precipitates with Spir-1, HEK293T cells were transfected with plasmids encoding Myc-tagged human Spir-1, Spir-2, β-TrCP, or Myc-GFP, together with plasmids expressing FLAG-tagged, codon-optimised K7 or another Bcl-2-like VACV protein, A49 (46). Immunoprecipitation with anti-Myc or anti-FLAG affinity resins showed that K7 was co-precipitated by Myc-Spir-1 (Fig 1A), but not by Myc-Spir-2, Myc-β-TrCP or Myc-GFP. In contrast, A49 interacted with β-TrCP as reported (47), but not with the other proteins. Reciprocally, FLAG-K7 co-precipitated Spir-1 (Fig 1B), but not the other Myc-tagged proteins, whilst β-TrCP only interacted with A49.

**Fig 1:**
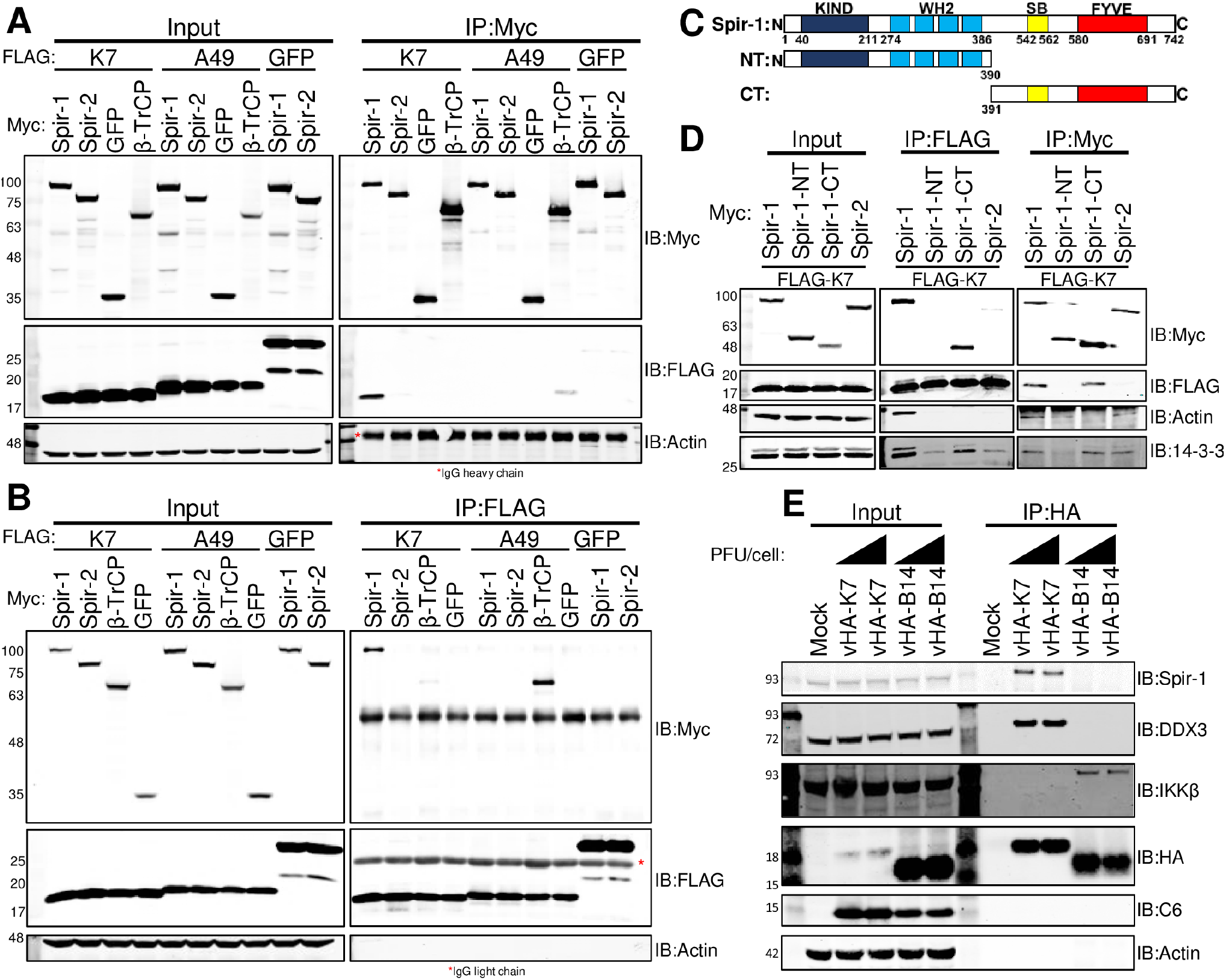
Spir-1 co-immunoprecipitates VACV protein K7 via its C-terminal region. HEK293T cells were transfected (**A, B** and **D**) with Myc-tagged and FLAG-tagged plasmids overnight. Cell lysates were immunoprecipitated using either Myc (**A** and **D** – right panel) or FLAG affinity resins (**B** and **D** – middle panel) and analysed by SDS-PAGE and immunoblotting. (**C**) Schematic representations of hSpir-1 isoform 2 full-length (top) and its C- and N-terminal truncations. (**E**) HEK293T cells were either mock-infected or infected at 5 or 10 PFU/cell with vHA-K7 or vHA-B14 for 4 h. Lysates were immunoprecipitated using HA-affinity resin and analysed by SDS-PAGE and immunoblotting. In (**A**), (**B**), (**D**) and (**E**) the positions of molecular mass markers in kDa are shown on the left. Each experiment was done 3 times and representative results are shown.

To investigate if the actin-binding domains of Spir-1 were needed for interaction with K7, two Myc-tagged truncations of Spir-1 were generated: an N-terminal region (amino acid residues 1-390), containing the four actin-binding WH2 domains and the KIND domain; and a C-terminal region, containing the SB and FYVE domains (amino acid residues 391-742, Fig 1C). These truncations, Spir-1 and Spir-2 were co-expressed with FLAG-K7 and Myc-tagged immunoprecipitation showed that the C terminus of Spir-1 was sufficient for interaction with K7, whereas the actin-binding domains were dispensable (Fig 1D, middle panel). The reciprocal IP gave the same conclusion and FLAG-K7 co-precipitated full-length Spir-1 and its C-terminal region (Fig 1D, right panel). As controls, the N-terminal region co-precipitated endogenous actin via its WH2 domains, and the C-terminal region interacted with 14-3-3, another binding partner of Spir-1 (48, 49).

To determine if Spir-1 and K7 interacted at endogenous levels, cells were either mock-infected or infected with VACV expressing HA-tagged K7 (vHA-K7) (10) or B14 (vHA-B14) (50). HA-immunoprecipitation confirmed that K7 interacts with endogenous Spir-1, whilst B14 does not (Fig 1E). As reported, K7 also co-precipitated with DDX3 (11), and B14 interacted with IKKβ (51). Altogether, these results indicate that Spir-1, but not Spir-2, interacts with K7 via its C-terminal region and independent of its actin-binding domains.

### Ectopic Spir-1 increases IRF3-dependent gene expression

Since the interaction of Spir-1 with K7 is independent of its actin-binding domains, and K7 is a VACV immunomodulator and virulence factor, we hypothesised that Spir-1 might have an unknown function in antiviral immunity. To test this, the impact of Spir-1 on innate immune signalling pathways was assessed by luciferase reporter gene assays. First, cells were transfected with a reporter plasmid in which the expression of firefly luciferase is driven by the IFNβ promoter. IFNβ expression was induced by Sendai virus (SeV) infection, which is sensed by RIG-I (52). Myc-Spir-1 expression alone did not affect IFNβ-dependent gene activation, however, Spir-1 augmented activation induced by SeV infection (Fig 2A). In contrast, Myc-tagged GFP had no effect, and VACV protein C6 was inhibitory as described (53). Spir-1 did not affect activation of ISRE-dependent gene expression downstream of type I IFN, whereas C6, but not N1, was inhibitory (54) (Fig 2B). Similarly, Spir-1 did not affect NF-κB activation in response to TNF-α, whilst B14 inhibited it (51) and VACV protein N2 had no effect (55) (Fig 2C). Further analysis using an IRF3-specific reporter plasmid (*ISG56*.*1* or *IFIT1* promoter) showed Spir-1 caused a dose-dependent increase in IRF3 activation induced by the CARD domain of RIG-I (Fig 2D, 2E). As controls, VACV protein N2 inhibited IRF3 activation as described (55) while A49, an NF-κB inhibitor (47), did not. Unlike Spir-1, Spir-2 did not enhance IRF3 activation whereas DDX3 did, as described (11) (Fig 2E).

**Fig 2:**
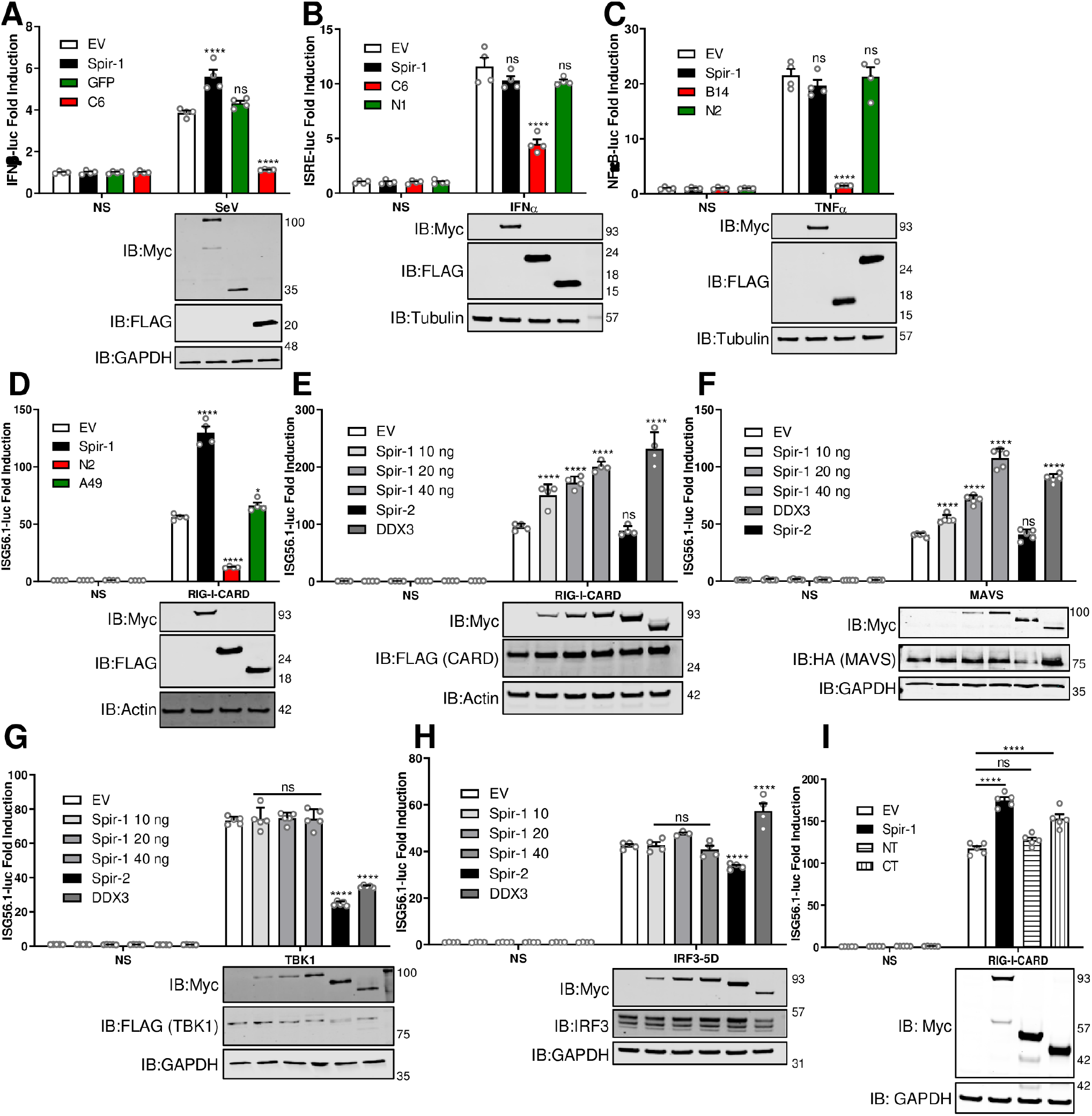
Ectopic expression of Spir-1 increases IRF3-dependent gene expression induced by other stimuli, at or downstream of MAVS. HEK293T cells were transfected with IFNβ (**A**), ISRE (**B**) and NF-κB (**C**) firefly luciferase reporter plasmids, together with TK-renilla luciferase and 40 ng of the plasmids for expression of the indicated proteins. After overnight transfection, cells were stimulated with SeV for 24 h (**A**), IFNα (**B**) or TNF-α (**C**) for 8 h. **D-I**) HEK293T cells were transfected with the ISG56.1 firefly luciferase reporter plasmid, TK-renilla luciferase and plasmids for expression of the indicated proteins. Cells were also co-transfected with EV as the non-stimulated (NS) controls or with the 5 ng of CARD-domain of RIG-I (**D, E** and **I**), 40 ng of MAVS (**F**), 40 ng of TBK-1 (**G**) and 5 ng of IRF3-5D (**H**) plasmids to activate the IRF3 pathway. EV was added to samples when necessary to keep the final amount of DNA transfected as 40 ng in all samples. Cell lysates were prepared and luciferase expression was measured and normalised to renilla luciferase. At least triplicate samples were analysed for each condition. Data are expressed as the mean (± SEM) fold induction of the firefly luciferase activity normalised to renilla values for the stimulated versus non-stimulated samples. Data shown are representative of three independent experiments. Immunoblots underneath each graph show the expression levels of the different proteins. The positions of molecular mass markers in kDa are shown on the right and the antibodies used are shown on the left. ns = not significant; *P < 0.05; ****P < 0.0001.

To map at which stage in the IRF3 pathway Spir-1 was acting, further *ISG56*.*1* reporter assays were performed in which the pathway was stimulated by expression of its different components. Spir-1 activated the IRF3 pathway in a dose-response manner when MAVS (mitochondrial antiviral-signalling protein) was used as stimulant (Fig 2F), but not when downstream components such as TBK1 (Fig 2G), IKKε (S1 Fig) or not when downstream components such or the constitutively active IRF3-5D (Fig 2H) were expressed. These findings indicate Spir-1 enhances IRF3 activation at or downstream of MAVS and upstream of IKKε or TBK1. Finally, N-and C-terminal fragments of Spir-1 were tested following stimulation by RIG-I CARD. Neither half of Spir-1 activated the pathway fully, but the C-terminal region had greater activity than the N-terminal region (Fig 2I).

### Spir-1 interaction with K7 is direct, independent of DDX3, and requires a diphenylalanine motif

K7 interacts directly with DDX3 (56, 57), an adaptor protein in the IRF3 pathway (58, 59). To investigate whether K7 binding to Spir-1 was via DDX3, cells stably expressing an inducible shRNA targeting DDX3 (shDDX3) (59) were used. Knockdown, rather than knockout, of DDX3 was utilised because *DDX3* encodes an essential protein (60). Cells expressing a non-silencing control (NSC) shRNA were used in parallel. DDX3 knockdown was induced by incubation with doxycycline (DOX) for 48 h as described (59), prior to transfection with either FLAG-tagged K7 or A49 for 24 h. Viral proteins were immunoprecipitated via their FLAG tag, followed by immunoblotting for endogenous Spir-1. DDX3 levels were reduced greatly in shDDX3 cells following DOX treatment, but not in control cells (Fig 3A). Despite this, Spir-1 co-precipitation by K7 was unaffected (Fig 3A). A49 did not co-precipitate either DDX3 or Spir-1 (Fig 3A). Next, the shDDX3 cells were used to determine if DDX3 contributed to Spir-1-induced IRF3 activation. After DDX3 knockdown, there was no significant difference in IRF3 stimulation by Spir-1 (Fig 3B). Notably, K7 inhibited IRF3 activation when DDX3 was knocked down, showing K7 had another IRF3 inhibitory mechanism independent of DDX3 (Fig 3B). Moreover, the presence of K7 reversed the activation of the pathway by Spir-1 (Fig 3B). In summary, Spir-1 interaction with K7 and its function in the IRF3 pathway are independent of DDX3.

**Fig 3:**
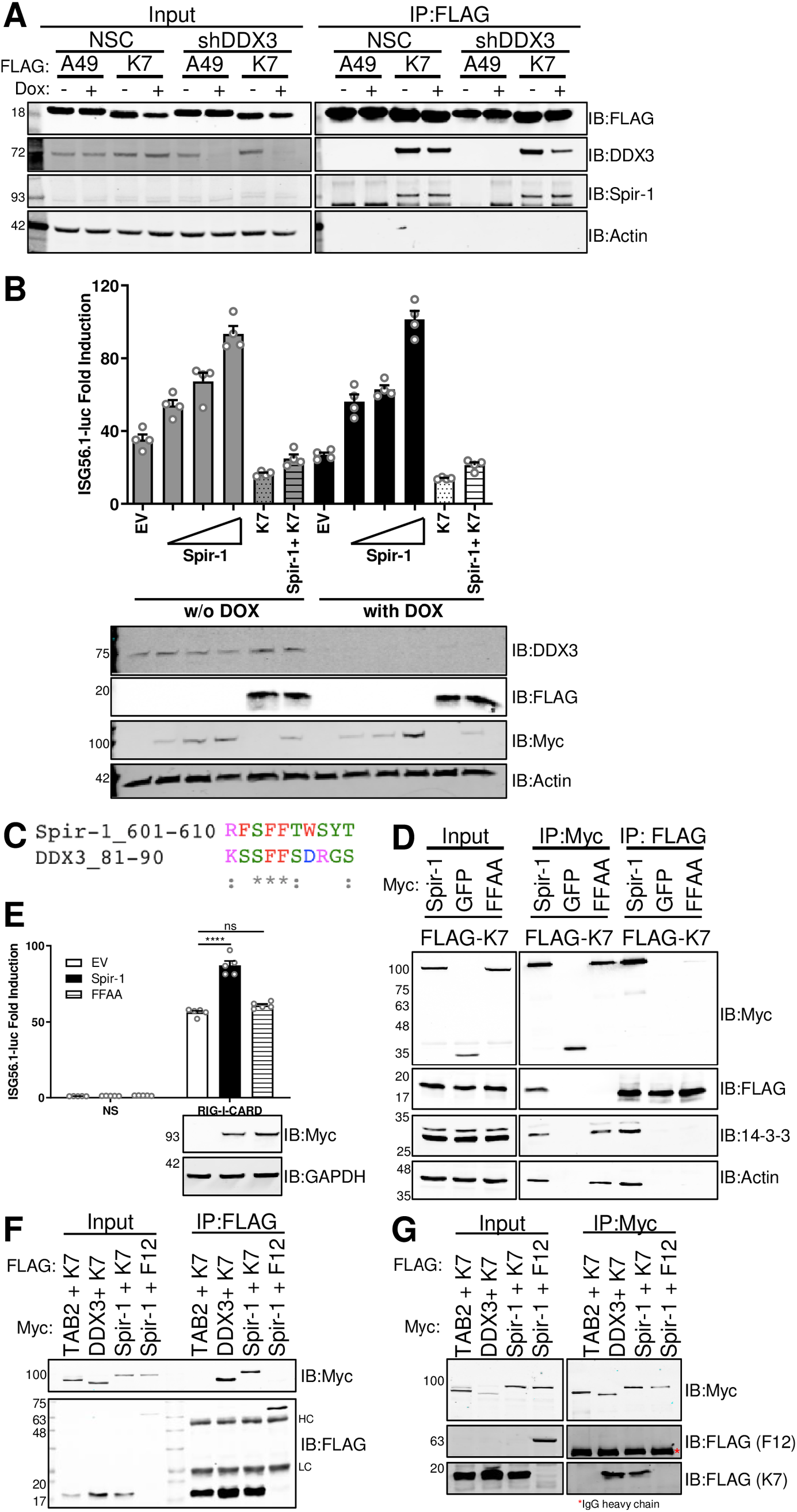
Spir-1 and DDX3 share a conserved diphenylalanine motif that is required for IRF3 activation and direct binding to K7. (**A**) DDX3 knockdown was induced in HEK293T cells stably transfected with pTRIPZ-shDDX3 (or a NSC pTRIPZ vector) through incubation with doxycycline for 48 h. Twenty-four h after doxycycline addition, cells were transfected with FLAG-tagged plasmids overnight. Cell lysates were immunoprecipitated using FLAG-affinity resin and analysed by SDS-PAGE and immunoblotting. (**B**) DDX3 knockdown was induced in HEK293T shDDX3 cells as in (**A**) and cells were then transfected with ISG56.1-firefly luciferase reporter, TK-renilla luciferase, plasmids for expression of the indicated proteins together with the CARD-domain. Cell lysates were prepared and analysed as in Fig 2. Immunoblots underneath the graph show the expression levels of the different proteins. (**C**) Alignment of amino acid residues of Spir-1 and DDX3 showing the conserved diphenylalanine motif. (**D**) HEK293T cells were transfected overnight with Myc-tagged Spir-1 wild type, GFP or Spir-1 mutant FFAA together with FLAG-K7. Cell lysates were immunoprecipitated using either Myc (middle panel) or FLAG-affinity resins (right panel) and analysed by SDS-PAGE and immunoblotting. (**E**) HEK293T cells were transfected with ISG56.1 firefly luciferase reporter, TK-renilla luciferase, plasmids for expression of the indicated proteins together with the CARD-domain of RIG-I or EV as the NS control. Cell lysates were prepared and analysed as in (**B**). The panel underneath the graph shows immunoblots for the expression level of K7 and GAPDH. (**F, G**) Myc or FLAG-tagged proteins were synthesized by *in vitro* transcription/translation. Samples were immunoprecipitated using FLAG-(**F**) or Myc-affinity resins (**G**) and analysed by SDS-PAGE and immunoblotting. For all immunoblots, the positions of molecular mass markers in kDa are shown on the left and the antibodies used on the right. ns = not significant; ****P < 0.0001. HC/LC: IgG heavy chain or light chain, respectively.

Since both DDX3 and Spir-1 bind to K7 and activate IRF3, their amino acid sequences were compared. The structure of K7 bound to a DDX3 peptide and subsequent structure-based mutagenesis showed that a diphenylalanine (FF) motif in DDX3 is essential for binding K7 and for IRF3 activation (57). An alignment of the C terminus of Spir-1 and the minimal ten-amino acid peptide from DDX3 (residues 81-90) needed for binding K7, showed a similar FF motif in Spir-1 (Fig 3C). To determine if this was important for Spir-1 binding to K7, the phenylalanines were mutated to alanines (FFAA) and the mutant was expressed in cells together with FLAG-tagged K7. Wild-type (WT) Myc-Spir-1 and Myc-GFP were also expressed. Anti-Myc or anti-FLAG immunoprecipitation showed that the FFAA mutation impaired Spir-1 interaction with K7 (Fig 3D), although it still precipitated both actin and 14-3-3, indicating normal folding. Notably, the FFAA mutant no longer activated IRF3 (Fig 3E).

To determine if the K7 interaction with Spir-1 was direct, these proteins were expressed by *in vitro* transcription and translation and immunoprecipitated alongside Myc-tagged TAB2 and FLAG-tagged VACV protein F12 (61) as controls. Both Spir-1 and DDX3 interacted directly with K7, whereas Spir-1 did not interact with F12, and K7 did not interact with TAB2 (Fig 3F, 3G). Taken together, this demonstrated that Spir-1 binds directly to K7 and shares with DDX3 a conserved diphenylalanine motif, which is required for its function in the IRF3 pathway.

### K7 uses the same amino acid residues to target both Spir-1 and DDX3

K7 binds to the DDX3 peptide through a hydrophobic pocket within a negatively-charged face (56, 57). Thus, interactions between K7 and the DDX3 peptide involve hydrophobic contacts, hydrogen bonds and electrostatic interactions, of which electrostatic contacts between R88 of DDX3 with both D28 and D31 of K7 are notable. D31 also forms a hydrogen bond with DDX3 S83 (57). Using this information, K7 mutants were generated in an attempt to distinguish between K7 binding to Spir-1 and DDX3. K7 D28 and D31 were changed singly or together to alanine (D28A, D31A) and expressed alongside WT K7 or GFP, and together with Myc-tagged Spir-1 and HA-tagged DDX3. Immunoprecipitation showed that K7 D28A still bound both Spir-1 and DDX3, whilst K7 D31A had impaired binding to both proteins, especially DDX3 (Fig 4A). Furthermore, the double mutant (DDAA) had further reduced binding to Spir-1 and no detectable binding to DDX3 (Fig 4A). Importantly, all mutants still interacted with COPε (Fig 4A), another K7 binding partner (13), suggesting normal K7 folding. Co-precipitation of endogenous Spir-1 with FLAG-tagged K7 and mutants gave similar conclusions (Fig 4B). Notably, the K7 D31A mutant inhibited IRF3 activation poorly compared to the WT or D28A (Fig 4C). Altogether, these data indicate that K7 requires D31 for interacting with both DDX3 and Spir-1 and via each interaction it inhibits the IRF3 pathway.

**Fig 4:**
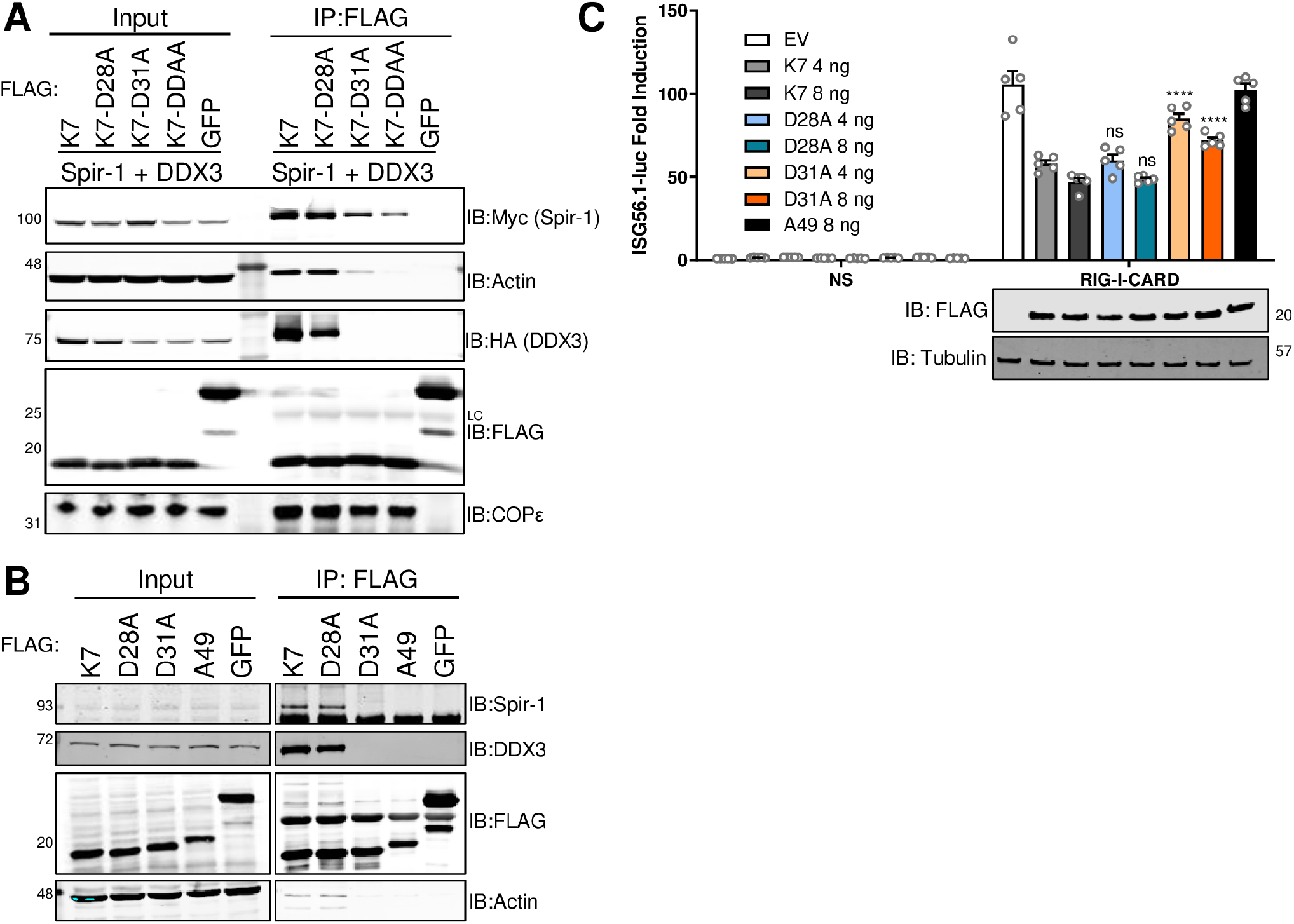
K7 residue Asp31 is important for binding to Spir-1 and DDX3 and inhibition of IRF3 activation. HEK293T cells were transfected with FLAG-tagged GFP, K7 wild type or mutants, Myc-Spir-1 and HA-DDX3 (**A**) or with only FLAG-tagged plasmids overnight (**B**). Cell lysates were immunoprecipitated using FLAG-affinity resin and analysed by SDS-PAGE and immunoblotting. The positions of molecular mass markers in kDa are shown on the left and the antibodies used on the right. (**C**) HEK293T cells were transfected with ISG56.1-firefly luciferase reporter, TK-renilla luciferase, plasmids for expression of the indicated proteins together with the CARD-domain or EV as the NS control. Cell lysates were prepared and analysed as in Fig 2. Statistical analyses compared the fold induction of the mutant sample to its respective K7 wild type. The panel underneath the graph shows immunoblots for the expression levels of the different K7 proteins and α-tubulin. The positions of molecular mass markers in kDa are shown on the right and the antibodies used on the left. ns = not significant; ****P < 0.0001. LC: IgG light chain.

### IRF3 activation is reduced in the absence of Spir-1

To investigate Spir-1-mediated activation of IRF3 further, Spir-1 knockout (KO) HEK293T cell lines were generated by CRISPR-Cas9-mediated targeting of *SPIRE1* exon 3, which is conserved in all Spir-1 isoforms (Fig 5A). After single cell selection, HEK293T cell lines were confirmed to lack Spir-1 by immunoblotting (Fig 5B) and by sequencing of both *Spire-1* alleles (Fig 5A). A clone that lacked Spir-1 protein expression and in which the *Spire-1* open reading frame was disrupted by frameshift mutations in both alleles, and that lacked WT sequence, was selected. Additionally, a vector expressing Myc-tagged Spir-1 was used to rescue Spir-1 expression in the KO cell line following lentiviral transduction. In parallel, both WT and KO cells were transduced with a control empty vector (EV) lentivirus. Spir-1 WT cells transduced with EV, and Spir-1 KO cells transduced with EV or Myc-Spir-1 were infected with SeV to activate the IRF3 pathway. Supernatants were collected for ELISA and cells lysates were used for either RNA extraction or immunoblotting, which confirmed that cells did or did not express Spir-1 (Fig 5B). The expression of Myc-Spir-1 increased in the presence of SeV, but this was not observed for endogenous Spir-1 (Fig 5B). One possible explanation is that Myc-Spir-1 expression is driven by the human cytomegalovirus immediate early promoter (62) that contains binding sites for transcription factors such as NF-κB (63), which can also be activated during SeV infection. The phosphorylation of IRF3 at S396 (p-IRF3), a hallmark of IRF3 activation (64), increased greatly 24 h after SeV infection of WT cells (Fig 5B) but was reduced in the absence of Spir-1 and restored in the KO cells expressing Myc-Spir-1 (Fig 5B). Moreover, after SEV infection, Spir-1 KO cells produced lower levels of mRNAs for *ISG56/IFIT1* (Fig 5C), *IFNB1* (Fig 5D) and *CXCL10* (Fig 5E), and secreted less CXCL-10 (Fig 5F), as measured by RT-qPCR and ELISA, respectively. Importantly, the absence of Spir-1 did not change the levels of CXCL-10 secreted in response to TNF-α, a specific agonist of the NF-κB pathway (Fig 5G).

**Fig 5:**
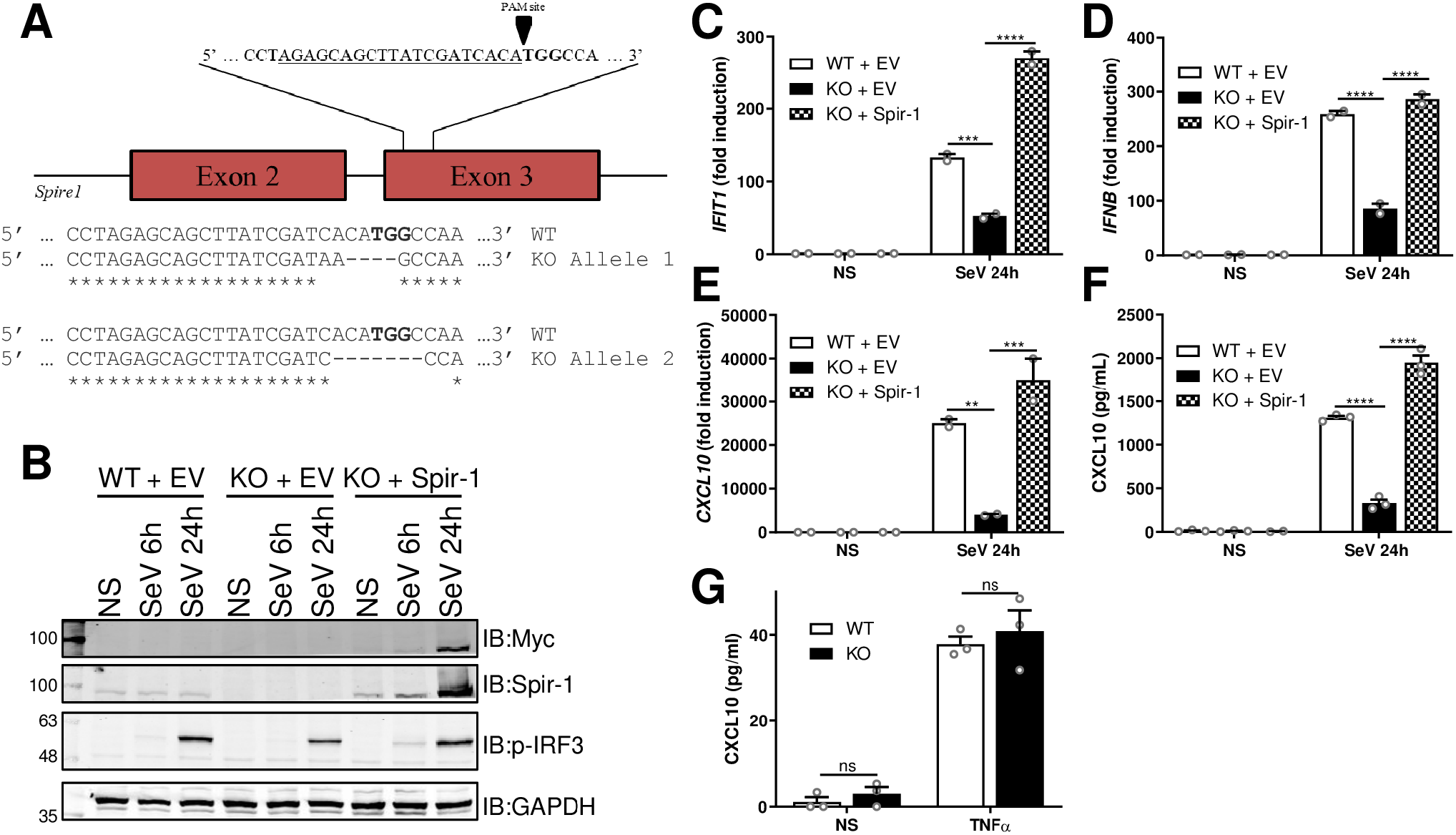
Spir-1 contributes to IRF3 phosphorylation and IRF3 stimulated gene expression after SeV infection in HEK293T cells. (**A**) Schematic of CRISPR-Cas9-mediated knockout strategy targeting *Spire1* exon 3 and single allele sequences of HEK293T Spir-1 knockout (KO) cells. (**B-F**) Spir-1 knockout (KO) HEK293T cells were transduced with empty vector (EV) or Myc-Spir-1 lentiviruses to rescue Spir-1 expression. Cells lines were either non-infected (NS) or infected with SeV for the indicated times. Cells were then either lysed and analysed by immunoblotting (**B**) or subjected to total RNA extraction followed by RT-qPCR (**C-E**). Supernatants were analysed by ELISA (**F**). (**G**) Spir-1 KO or wild type (WT) HEK293T cells were stimulated overnight with TNFα and supernatants were analysed by ELISA. ns = not significant, **P < 0.01, ***P < 0.001, ****P < 0.0001.

Immortalised mouse embryonic fibroblasts (MEFs) from mice with a terminator (gene trap) between exons 3 and 4 of the *Spire1* gene (65) were used to investigate the contribution of Spir-1 to IRF3 activation in cells from a different mammal. After stimulation of IRF3 by either poly I:C or SeV infection, MEFs were lysed for immunoblotting or RNA analysis and supernatants were collected for ELISA. A reduction in p-IRF3 was seen in Spir-1 KO cells (Fig 6A, 6B) and this was quantified (bottom graphs in Fig 6A, 6B). The Spir-1 KO MEFs also expressed less mRNAs for cytokines and chemokines upon stimulation with poly I:C or SeV infection than their WT controls (Fig 6C, 6D, 6E, 6F, 6G, 6H). Likewise, CXCL-10 and IL-6 secretion were lower from Spir-1 KO cells (Fig 6I, 6J, 6K, 6L). However, the levels of CXCL-10 and IL-6 secreted after stimulation with IL-1β were not reduced in Spir-1 KO MEFs (Fig 6M, 6N); indeed, there was a small increase in IL-6 levels in KO cells (Fig 6N). In summary, in both mouse and human cells lacking Spir-1 there is a defect in IRF3 activation downstream of RIG-I/MDA5, whilst other immune signalling pathways were largely unaltered.

**Fig 6:**
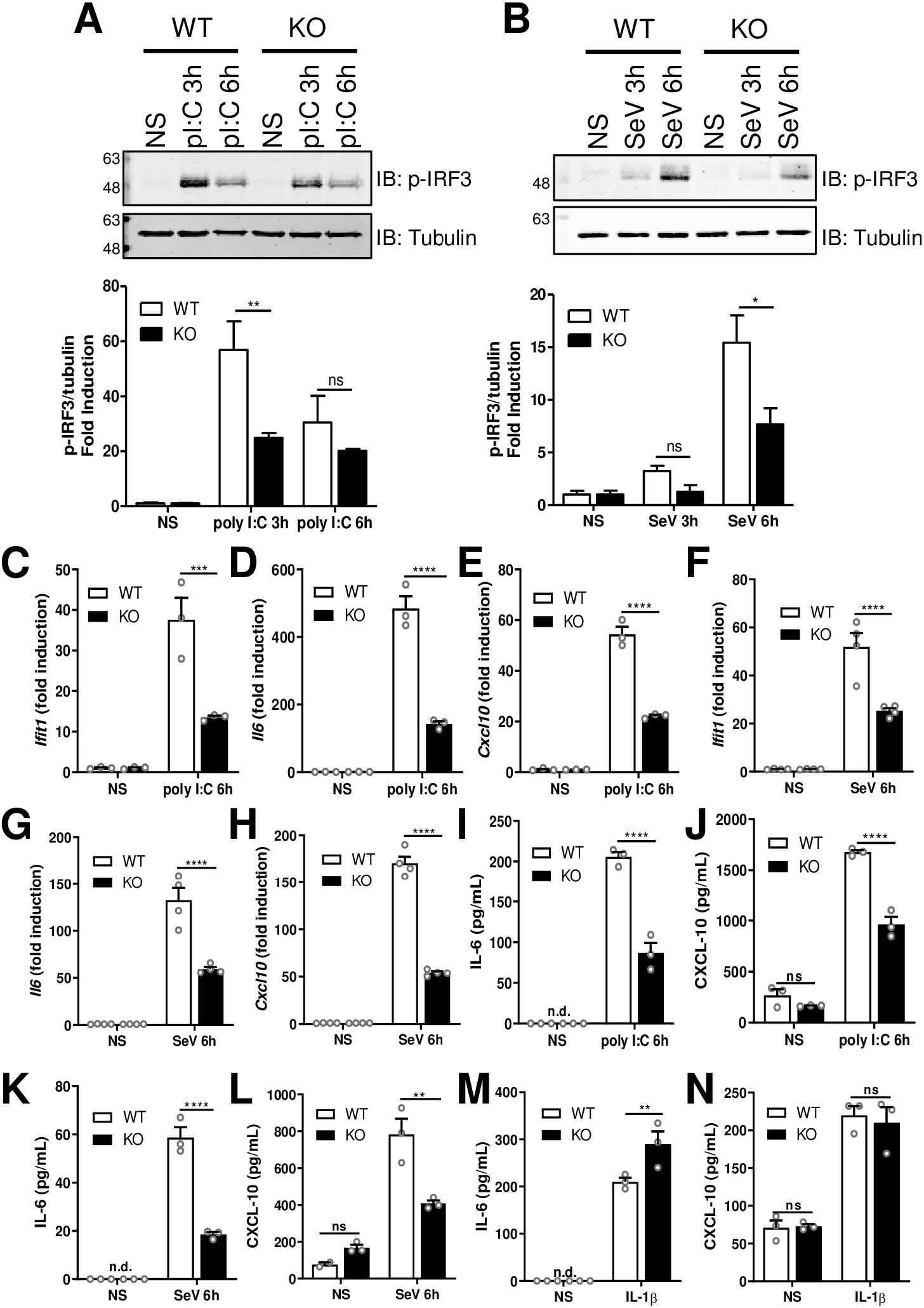
IRF3 activation is reduced in Spir-1 KO MEFs. Spir-1 WT or KO MEF cells were transfected with poly I:C (**A, C-E, I-J**) or infected with SeV (**B, F-H, K-L**) for the indicated times. Cells were lysed and analysed by immunoblotting (**A, B**) and phospho-IRF3 bands intensity was quantified and normalised by the intensity of α-tubulin from at least two different experiments (**A, B**, bottom graphs). The positions of molecular mass markers in kDa are shown on the left and the antibodies used on the right. Cells were also subjected to total RNA extraction followed by RT-qPCR (C-H) and supernatants were analysed by ELISA (**I-L**). (**M, N**) Spir-1 WT or KO MEF cells were stimulated with IL-1β for 8 h and supernatants were analysed by ELISA. ns = not significant, *P < 0.05, **P < 0.01, ****P < 0.0001.

### Spir-1 is a viral restriction factor

Given that Spir-1 interacts with VACV protein K7 and enhances IRF3 activation, which is critical for the host response to viral infection, the impact of Spir-1 on VACV replication and spread was assessed. Spir-1 WT and KO HEK293T cells and the derivative cell line in which Spir-1 expression was restored in the KO cells, were infected with A5-GFP-VACV, a VACV strain expressing GFP fused to capsid protein A5 (66), and 2 days (d) later plaque diameters were measured (Fig 7A, 7B). A significant increase in plaque size was seen in the absence of Spir-1 compared to both WT and rescued cells (Fig 7B), suggesting that Spir-1 restricts VACV spread and/or replication. Lastly, virus growth on those cells was assessed. Cells were infected with 0.001 plaque-forming unit (PFU)/cell of VACV strain WR and 2 d later the virus yield was determined by plaque assay. Spir-1 KO cells yielded higher viral titres compared to both WT and Spir-1-complemented cells (Fig 7C).

**Fig 7:**
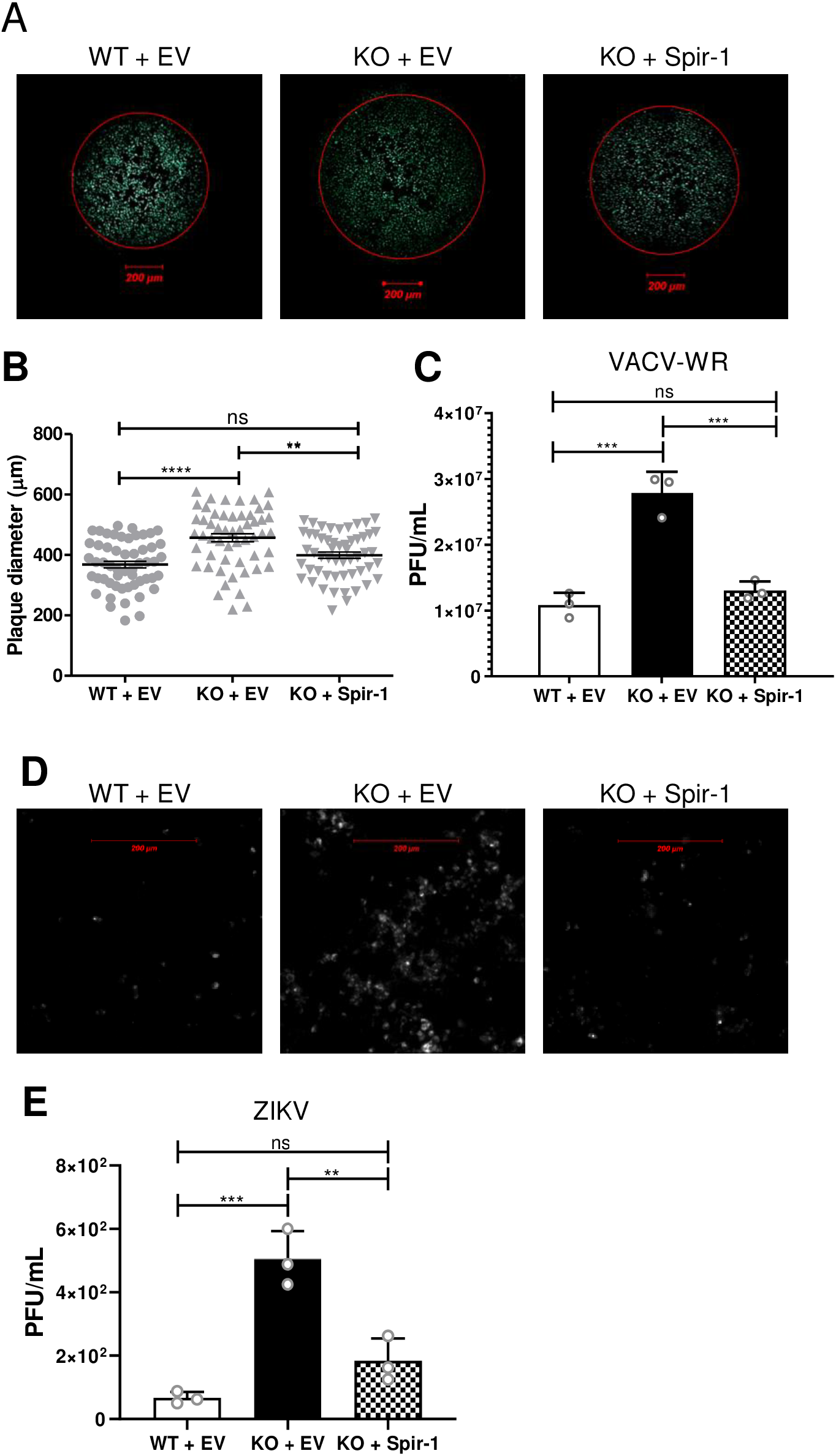
Spir-1 is a cellular restriction factor for VACV and ZIKV. (**A, B**) Spir-1 WT or KO and Spir-1 KO complemented HEK293T cells were infected with VACV-A5-GFP and plaque diameters were measured at 48 h p.i. (**A**) Representative plaques formed in each cell line. (**B**) Plaques diameter measurements (n=54). (**C**) Spir-1 WT or KO and Spir-1 KO complemented HEK293T cells were infected with VACV WR at 0.001 PFU/cell for 48 h and the virus yield was measured in BSC-1 cells. ns = not significant, *P<0.05, **P < 0.01, ***P < 0.001, ****P < 0.0001. (**D and E)** Spir-1 WT or KO and Spir-1 KO complemented HEK293T cells were infected with ZIKV-mCherry at 0.01 PFU/cell for 72 h and ZIKV-infected monolayers were imaged (**D**) or the virus yield was measured in VERO E6 cells (**E**). ns = not significant, **P < 0.01, ***P < 0.001.

The IRF3 pathway also restrict RNA viruses, therefore the impact of Spir-1 in restricting Zika virus (ZIKV), a single-stranded RNA virus, was also assessed. HEK293T Spir-1 WT, KO and Spir-1-rescued cells were infected with ZIKV-mCherry with 0.01 PFU/cell. Three days later, the mCherry signal was visualised by live-cell imaging and showed a greater signal in the KO cells compared to controls (Fig 7D), suggesting an increased ZIKV replication in these cells. To address that possibility, the virus yield in the supernatant of infected HEK293T cells was determined by plaque assay in VERO E6 cells (Fig 7E). In the absence of Spir-1, there is an increase in ZIKV titres compared to the WT and Spir-1 restored KO cells. Collectively, loss of Spir-1 caused an increase in both virus plaque size and yield of infectious virus, showing that Spir-1 functions as a restriction factor against both a DNA (VACV) and an RNA (ZIKV) virus.

## Discussion

During co-evolution with their hosts, viruses have evolved mechanisms to evade the host antiviral responses and thereby replicate and spread efficiently. Such virus proteins are numerous and have diverse functions and, therefore, can be used to gain insight into how the host innate immune system functions. VACV is a poxvirus and encodes scores of immunomodulatory proteins many of which target the innate immune system. In several cases, multiple VACV proteins engage the same pathway, but despite this, these proteins each contributes to virulence indicating non-redundant functions (9). Sometimes a single protein can inhibit multiple pathways (9) or an open reading frame can encode multiple proteins with different functions (67). High-throughput proteomic approaches have been used to identify binding partners of VACV proteins, or cellular proteins that are up or down-regulated during virus infection, and such cellular proteins may have antiviral activity and function in innate immunity (14, 16). For example, histone deacetylases (HDAC) 4 and 5 are each targeted for proteasomal degradation by VACV protein C6 and function as a virus restriction factor (15, 16).

Here, Spir-1 is characterised as a new cellular protein that is bound by VACV protein K7 (Fig 1), a virulence factor that inhibits NF-κB and IRF3 activation (10, 11). K7 binding to Spir-1 was identified in a proteomic screen for binding partners of virus immunomodulatory proteins (14). Here, this interaction is confirmed, demonstrated to occur at endogenous levels during VACV infection (Fig 1E), and shown to be direct (Fig 3F, 3G).

Spir-1 belongs to a family of proteins involved in actin organisation (44) but K7 interaction with Spir-1 does not require its N-terminal actin-binding domain (Fig 1D). VACV induces polymerisation of actin to facilitate virus dissemination from the cell surface and VACV mutants unable to polymerise actin spread poorly and form small plaques (68). However, a mutant lacking K7 produces normal size plaques (10). Given that K7 is an immunomodulatory protein, we hypothesised that Spir-1 might have a function in innate immunity. In reporter gene assays, Spir-1 over-expression enhanced IRF3-dependent gene expression (Fig 2D) but did not affect JAK-STAT signalling induced by type I IFN, or NF-κB activation (Fig 2B, 2C). Conversely, human and murine Spir-1 knockout cell lines showed diminished IRF3 signalling, with reduced IRF3 phosphorylation and reduced transcription and expression of IRF3-responsive genes (Fig 5B, 6A, 6B). Importantly, rescue of Spir-1 expression in Spir-1 KO cells restored phospho-IRF3 and cytokine expression levels (Fig 5).

To map where Spir-1 influences IRF3 activation, the pathway was activated by different agonists and the influence of Spir-1 examined. Spir-1 enhanced activation induced by SeV infection (Fig 2A, 5 and 6), poly I:C transfection (Fig 6), and overexpression of both RIG-I CARD (Fig 2D, 2E) and MAVS (Fig 2F), but not overexpression of TBK1, IKKε or a constitutively active IRF3 (Fig 2G and S1). SeV RNA is sensed by RIG-I (69) whilst high molecular weight (HMW) poly I:C is sensed by MDA-5 (52, 70). This suggests Spir-1 acts at or downstream of MAVS, the adaptor molecule recruited downstream of RIG-I and MDA5 activation (71). The association of RIG-I/MDA5 with MAVS results in the recruitment of tumour necrosis factor receptor-associated factors (TRAF), such as TRAF3, leading to the phosphorylation of IRF3 by TBK1/IKKε. Collectively, these data suggest Spir-1 functions at MAVS or TRAF3 level.

K7 inhibition of IRF3 activation was attributed to direct binding to DDX3, which acts as a multifunctional scaffolding adaptor culminating in IRF3 activation (58, 59). However, data presented here show K7 has additional targets within the IRF3 pathway, since Spir-1 also contributes to IRF3 activation and K7 is able to inhibit IRF3 activation even in the absence of DDX3. Furthermore, DDX3 and Spir-1 act at different positions in the pathway; DDX3 acts at the TRAF3 and TBK1/IKKε level (9, 11, 59, 72), whilst Spir-1 acts upstream of TBK1/IKKε (Fig 2). Interestingly, both DDX3 and Spir-1 share a diphenylalanine motif that is responsible for both binding to K7 and their function in the IRF3 pathway (Fig 3) (57). Furthermore, the same amino acid residue in K7 (D31) is crucial for its binding to both cellular proteins (Fig 4). This prevented assessment of the relative importance of K7 binding to each cellular protein.

Nonetheless, Spir-1 has an important antiviral role because cells lacking Spir-1 show enhanced VACV and ZIKV replication and/or spread (Fig 7). Comparable data for DDX3 are lacking, and obtaining such data is complicated by DDX3 being an essential protein (60). Both K7 and Spir-1 localise in the cytoplasm (10, 34). Interestingly, mutations in the FYVE domain and in the diphenylalanine motif in Spir-1 change its sub-cellular localisation from a trans-Golgi network and post-Golgi vesicles localisation to an even cytoplasmic distribution (29, 34). Neither the N- or C-terminal halves of Spir-1 were capable of activating the IRF3 pathway as well as full-length protein (Fig 2I), despite comparable expression levels, suggesting that both halves are important. It is possible that Spir-1 is directed to the correct location by its C-terminal domain, whist its N-terminal domain mediates Spir-1 engagement with other proteins to promote IRF3 activation. This might explain why the FFAA Spir-1 mutant is no longer able to activate the IRF3 pathway. Another possibility is that K7 may bind to the Spir-1 diphenylalanine motif and displace other proteins or sequester it and prevent it reaching its subcellular location. Spir-1 has several isoforms (45), and one, Spire-1C (also known as Spire1-E13), differs from the canonical isoform by encoding an extra exon sequence (ExonC or E13), which mediates its mitochondrial localisation and regulates mitochondrial division (41). Even though this was not the isoform used in the present study, K7 might regulate Spir-1 trafficking to the mitochondrion, where MAVS is anchored (73).

RIG-I and MDA5 can sense RNA virus genomes directly, but also RNA generated by infection with DNA viruses (74), leading to IRF3 activation. Therefore, RIG-I/MDA5-triggered antiviral response is antagonised by several RNA and DNA viruses (71, 75). For instance, ZIKV has been shown to be sensed by both RIG-I and MDA-5 depending on the infected cell type (reviewed by (76)). Furthermore, it has long been known that dsRNA is generated during VACV infection (77-79), so it is not surprising that VACV encodes proteins to interfere with cytosolic RNA sensing. For instance, VACV protein E3 binds to dsRNA and prevents RIG-I-dependent sensing of RNA products generated via the transcription of AT-rich DNA by RNA polymerase III (80, 81), and the decapping enzymes D9 and D10 prevent the accumulation of dsRNA from viral complementary RNA molecules (82). Downstream of RNA sensing, VACV encodes proteins to inhibit IRF3 activation, such as C6, which acts at TBK1/IKKε activation level (53), N2, which acts downstream of IRF3 translocation to the nucleus (55) and K7, which targets DDX3 (11). Here, we show that Spir-1 also contributes to IRF3 activation, with both Spir-1 and DDX3 sharing a similar motif which binds K7.

In summary, Spir-1 is a new cellular protein that affects IRF3 activation and is a virus restriction factor. By exploring in detail the function of VACV protein K7, an additional cellular factor that regulates IRF3 activation and acts as a viral restriction factor has been characterised.

## Material and Methods

### Cells lines

Human embryonic kidney (HEK) 293T, HEK293T NSC or shDDX3 (kind gifts from Dr. Martina Schröder, Maynooth University, Ireland), murine embryonic fibroblasts (MEFs) Spir-1 KO and WT (kindly provided by Prof. Dr. Eugen Kerkhoff - University Hospital Regensburg, Germany), BSC-1 and VERO E6 (African Green Monkey) cells were cultured in Dulbecco’s modified Eagle’s medium (DMEM, Gibco) supplemented with 10% heat-treated (56 °C, 1 h) foetal bovine serum (FBS, Pan Biotech), 100 U/mL penicillin and 100 μg/mL streptomycin (P/S, Gibco). RK13 cells were grown in minimal essential medium (MEM, Gibco) supplemented with 10% FBS and P/S.

### Viruses

VACV strain Western Reserve (WR) and derivative strains expressing GFP fused to the capsid protein A5 (A5-GFP-VACV) (66), VACV expressing HA-tagged B14 (50) and VACV expressing HA-tagged K7 (10) were described. VACV strains were grown on RK13 cells and titrated by plaque assay on BSC-1 cells. Sendai virus (SeV) Cantell strain (Licence No. ITIMP17.0612A) was a gift from Prof. Steve Goodbourn (St George’s Hospital Medical School, University of London). ZIKV engineered to express a mCherry marker (83) was a kind gift from Dr. Trevor Sweeney (Department of Pathology, University of Cambridge).

### Plasmids

A plasmid encoding the Myc-tagged human Spir-1 isoform 2 and human Spir-2 (20) constructs were a kind gift of Prof. Dr. Eugen Kerkhoff. Spir-1 N- and C-terminal truncations were constructed by PCR amplification from pcDNA3.1-Myc-Spir-1 and inserted into plasmid pcDNA3.1-Myc. Codon-optimised FLAG-K7 was described (84) and Spir-1 and K7 plasmids were used as templates for site-directed mutagenesis according to the instructions of the QuikChange Site-Directed Mutagenesis Kit (Agilent). All mutations were confirmed by DNA sequencing. Myc-DDX3 and HA-DDX3 (11) were kindly provided by Dr. Martina Schröder. The IFNβ-firefly luciferase reporter plasmid was from T. Taniguchi (University of Tokyo, Japan), NF-κB-firefly luciferase was from R. Hofmeister (University of Regensburg, Germany) and ISG56.1-firefly luciferase was a gift from Ganes Sen (Cleveland Clinic, USA). ISRE-firefly luciferase and pTK-renilla luciferase (pRL-TK) plasmids were from Promega. Vectors expressing MAVS, IKKε, TBK1 and IRF3-5D (53), and RIG-I-CARD domain (85) were described. Lentivirus vector plasmid pLKO.DCMV.TetO.mcs (pLDT) (86) was a gift from Prof. Roger Everett (University of Glasgow, UK) and was used as backbone for sub-cloning the Myc-Spir-1. Plasmids pCMV.dR8.91 (expressing all necessary lentivirus helper functions) and pMD-G (expressing the vesicular stomatitis virus envelope protein G) were from Dr. Heike Laman (University of Cambridge, UK). pF3A WG plasmid was from Promega. px459 CRISPR-Cas9 plasmid was purchased from Addgene. More information about plasmids and oligonucleotide primers used are given in the reagents table.

### Antibodies and Reagents

Primary antibodies used were from the following sources: rabbit (Rb) anti-Myc (Cell Signaling, 2278), mouse (Ms) anti-Myc (Cell Signaling, 9B11), Rb anti-Actin (Sigma, A2066), Ms anti-FLAG (Sigma F1804), Rb anti-14-3-3 (Santa Cruz, sc-629), Ms anti -Spir-1 (Santa Cruz, sc-517039), Ms anti-Spir-1 (Abcam, ab57463), Rb anti-DDX3 (Cell Signaling, 2635), Rb anti-IKKβ (Cell Signaling, 2684), Rb anti-HA (Sigma, H6908), Ms anti-α-Tubulin (Millipore, 05-829), Ms anti-GAPDH (Sigma, G8795), Rb anti-IRF3 (Cell Signaling, 4962), Ms anti-COPε (Santa Cruz, sc-133194), Rb anti-phospho-IRF3 Ser396 (Cell Signaling, 4947S) and Rb polyclonal anti-C6 (53). For dilutions used for the primary antibodies, see reagents table. Secondary antibodies used (1:10,000 dilution) were IRDye 680RD-conjugated goat anti-rabbit IgG or anti-mouse IgG and IRDye 800CW-conjugated goat anti-rabbit IgG or anti-mouse IgG (LI-COR).

Reagents used in this study were: Anti-c-Myc Agarose from Santa Cruz Biotechnology, and monoclonal Anti-HA-Agarose, clone HA-7, ANTI-FLAG M2 Affinity Gel and Poly-D-lysine hydrobromide (all from Sigma Aldrich). Human IFNα, human TNF-α and mouse IL-1β were from Peprotech, HMW poly(I:C) and puromycin were from InvivoGen, and doxycycline was from Melford.

### Reporter Gene Assay

HEK293T cells were seeded in 96-well plates with 1.5 × 10^4^ cells per well. After two days, cells were transfected with 60 ng per well of the firefly luciferase reporter plasmids (IFNβ, ISRE, NF-κB or ISG56.1), 10 ng per well of pTK-renilla luciferase and different amounts of the expression plasmid under test or empty vector (EV) control using polyethylenimine (PEI, CellnTec, 2 μL per 1 μg DNA). When necessary, EV plasmid was added to the transfection so that the final amount of DNA transfected was kept constant. In cases where stimulation was done by transfecting another plasmid, the same amount of EV was transfected to the non-stimulated (NS) control wells. Cells were stimulated as shown in the Figures: (i) infection with SeV for 24 h (IFNβ Luc), (ii) 10 ng/mL of TNF-α for 8 h (NF-κB Luc) or (iii) with 1000 U/mL of IFNα for 8 h (ISRE-Luc). After stimulation, cells were washed with PBS, lysed with 100 uL/well of passive lysis buffer (Promega) and firefly and renilla luciferase activities were measured using a FLUOstar luminometer (BMG). The firefly luciferase activity in each sample was normalised to the renilla luciferase activity and fold inductions were calculated relative to the non-stimulated controls for each plasmid. In all cases, data shown are representative from at least three independent experiments with at least triplicate samples analysed for each condition.

### Immunoblotting

HEK293T (8 × 10^5^) or MEFs (2 × 10^5^) cells were seeded in 6-well plates and 24 h later cells were stimulated by infection with SeV or transfection with 5 µg/mL of poly I:C using lipofectamine 2000 (Life Technologies). After stimulation, cells were washed twice with ice-cold PBS, and scrapped into a cell lysis buffer containing 50 mM Tris-HCl pH 8, 150 mM NaCl, 1 mM EDTA, 10% (v/v) glycerol, 1% (v/v) Triton X-100 and 0.05% (v/v) NP-40, supplemented with protease (cOmplete Mini, Roche) and phosphatase inhibitors (PhosSTOP, Roche). Protein concentration was determined using a bicinchoninic acid protein assay kit (Pierce) before being boiled at 100 °C for 5 min. Proteins were then separated by SDS-polyacrylamide gel electrophoresis and transferred onto a nitrocellulose Amersham Protran membrane (GE Healthcare). Membranes were blocked at room temperature with either 5% (w/v) milk or 5% (w/v) bovine serum albumin (BSA, Sigma) in PBS containing 0.1% Tween 20. Then, membranes were incubated with a specific primary antibody diluted in blocking buffer at 4 °C overnight. After washing, membranes were probed with LI-COR secondary antibodies at room temperature followed by imaging using the LI-COR Odyssey imaging system, according to the manufacturer’s instructions. Where indicated, protein bands from at least two independent experiments were quantified by using Odyssey software (LI-COR Biosciences).

### Immunoprecipitation

HEK293T cells were seeded in 10-cm dishes (3.5 × 10^6^ cells per dish) and transfected with the plasmids indicated in the Figures using PEI. The following day, cells were washed twice with PSB, lysed in immunoprecipitation (IP) lysis buffer (150 mM NaCl, 50 mM Tris-HCl pH 7.4, 0.5% (v/v) Nonidet P-40 (NP-40) and protease (cOmplete Mini, Roche) and phosphatase inhibitors (PhosSTOP, Roche)) and cleared by centrifugation at 21,000 *g* for 15 min at 4 °C. Cleared lysates were then incubated with 20 μL of ANTI-FLAG M2 Affinity Gel (Sigma Aldrich) or Anti-HA Agarose (Sigma Aldrich) for 2 h, or with 50 uL of Anti-c-Myc Agarose overnight, at 4 °C. Alternatively, proteins were expressed by *in vitro* transcription/translation using the TNT SP6 High-Yield Wheat Germ Protein Expression System (Promega) prior to incubation with the affinity resins. Immunoprecipitations were washed 3 or 4 times in IP buffer and bound proteins were eluted in Laemmli SDS-PAGE loading buffer and heated at 100 °C for 5 min. Samples were then analysed by SDS-PAGE and immunoblotting.

### RT-qPCR

HEK293T (4 × 10^5^) or MEFs (1 × 10^5^) cells were seeded in 12-well plates. The next day, cells were stimulated by infection with SeV or transfection with 5 µg/mL of poly I:C using lipofectamine 2000 (Life Technologies). RNA was extracted using the RNeasy kit (QIAGEN) and 500 ng of each RNA sample was used to synthesise cDNA using Superscript III reverse transcriptase according to the manufacturer’s protocol (Invitrogen). mRNA was quantified by real-time PCR using a ViiA 7 Real-Time PCR System (Life Technologies), fast SYBR Green Master Mix (Applied Biosystems) and the following primers: human *CXCL10* (Fwd: GTGGCATTCAAGGAGTACCTC, Rev: GCCTTCGATTCTGGATTCAGA), human *IFNB1* (Fwd:ACATCCCTGAGGAGATTAAGCA, Rev: GCCAGGAGGTTCTCAACAATAG), human *IFIT1* (Fwd: CCTGAAAGGCCAGAATGAGG, Rev: TCCACCTTGTCCAGGTAAGT), human *GAPDH* (Fwd: ACCCAGAAGACTGTGGATGG, Rev: TTCTAGACGGCAGGTCAGGT), mouse *Ifit1* (Fwd: ACCATGGGAGAGAATGCTGAT, Rev: GCCAGGAGGTTGTGC), mouse *Il6* (Fwd: GTAGCTATGGTACTCCAGAAGAC. Rev: ACGATGATGCACTTGCAGAA), mouse *Cxcl10* (Fwd: ACTGCATCCATATCGATGAC, Rev: TTCATCGTGGCAATGATCTC) and mouse *Gapdh* (Fwd: ATGGTGAAGGTCGGTGTGAACGG, Rev: TTACTCCTTGGAGGCCATGTAGGC). Gene amplification was normalised to *GAPDH* (glyceraldehyde-3-phosphate dehydrogenase) amplification from the same sample, and the fold induction of genes in stimulated samples was calculated relative to the unstimulated control. Experiments were performed in at least biological duplicate and conducted at least twice.

### ELISA

HEK293T (4 × 10^5^) or MEFs (1 × 10^5^) cells were seeded in 12-well plates. The following day, cells were stimulated by: (i) infection with SeV; (ii) transfection with 5 µg/mL of poly I:C using lipofectamine 2000 (Life Technologies), (iii) 50 ng/mL of human TNF-α or (iv) 50 ng/mL of mouse IL-1β. After stimulation, supernatants were assayed for human or murine CXCL-10 and IL-6 protein using Duoset enzyme-linked immunosorbent assay (ELISA) reagents (R&D Biosystems) according to the manufacturer’s instructions.

### CRISPR/Cas9-mediated genome editing

Two guide RNAs (gRNAs) were designed using online software (http://tools.genome-engineering.org) to target *SPIRE1* gene exon 3, which is shared by all Spir-1 isoforms. CRISPR/Cas9-mediated genome editing of HEK293T cells was performed as described (87). Briefly, px459 CRISPR/Cas9 plasmids with or without gRNA sequence were transfected into HEK293T using TransIT-LT1 transfection reagent (Mirus, MIR 2306). Puromycin (1 μg/mL) was added to transfected cells and, after 48 h, puromycin-resistant cells were serially diluted to obtain individual clones. Several clones were amplified, and a few potential knockout clones were selected by immunoblotting and confirmed by genomic DNA sequencing at the gRNA target sites. Only one gRNA was successful and one clonal cell line was confirmed to be knockout after PCR-amplified genomic DNA was cloned into bacterial plasmids and multiple colonies (n = 20) were sequenced. These all contained frameshift mutations. No wild type allele was identified.

### Lentivirus transductions

Lentivirus particles for transduction were generated after transient co-transfection of HEK293T cells seeded in 6-cm dishes. Cells were transfected with pCMV.dR8.91 and pMD-G vectors together with either pLDT-EV or pLDT-Myc-Spir-1. After 48 h and 72 h, the supernatant was collected and passed through a 0.45 μm filter. Lentivirus-containing supernatant was then used to infect HEK293T Spir-1 WT and KO cells. Transduced cells were selected with 1 μg/mL puromycin followed by clonal selection by serial dilution. Spir-1 expression was assessed by immunoblotting.

### Virus Infection

Plaque size analysis were performed in HEK293T Spir-1 WT and KO cells seeded in 6-well plates coated with poly-D-lysine (Sigma). Once confluent, cells were infected with VACV-A5-GFP at 20 PFU per well for 2 d. Virus plaques diameters (*n*=54) were measured using AxioVision 4.8 software and a ZEISS Axio Vert.A1 fluorescent microscope.

Viral replication was measured by multi-step growth analyses of HEK293T Spir-1 WT and KO cells infected with VACV-WR at 0.001 PFU / cell or ZIKV-mCherry at 0.01 PFU/cell. For VACV, at 48 h post infection, cells were scraped in their medium and collected by centrifugation at 500 *g* for 5 min. Cells were subjected to three rounds of freeze-thawing before the infectious viral titre was determined by plaque assay on BSC-1 cells for 3 days. For ZIKV, at 72 h post infection, supernatants of infected cells were collected, and virus infectivity was determined by plaque assay on Vero E6 cells for 5 days. ZIKV-infected monolayers were also imaged using a Zeiss Axiovert 200M microscope. Images in Figure 7D were processed using Adobe Photoshop 2020 to enhance linearly the mCherry visualisation. SC-1 and VERO E6 cells were then fixed with 4% paraformaldehyde (PFA) and stained with toluidine blue.

### Statistical Analysis

Statistical analysis was carried out using one or two-way ANOVA test where appropriate with the Bonferroni post-test, using the GraphPad Prism statistical software (Graph-Pad Software). Statistical significance is expressed as follows: ns = not significant, *P < 0.05, **P < 0.01, ***P < 0.001, ****P < 0.0001.

## Acknowledgements

We thank Dr. Martina Schröder, (Maynooth University, Ireland) and Prof. Dr. Eugen Kerkhoff (University Hospital Regensburg, Germany) for providing cells lines and plasmids for this study. We also thank Callum Talbot-Cooper for critical reading of the manuscript.

## Supporting information

**S1 Fig.**
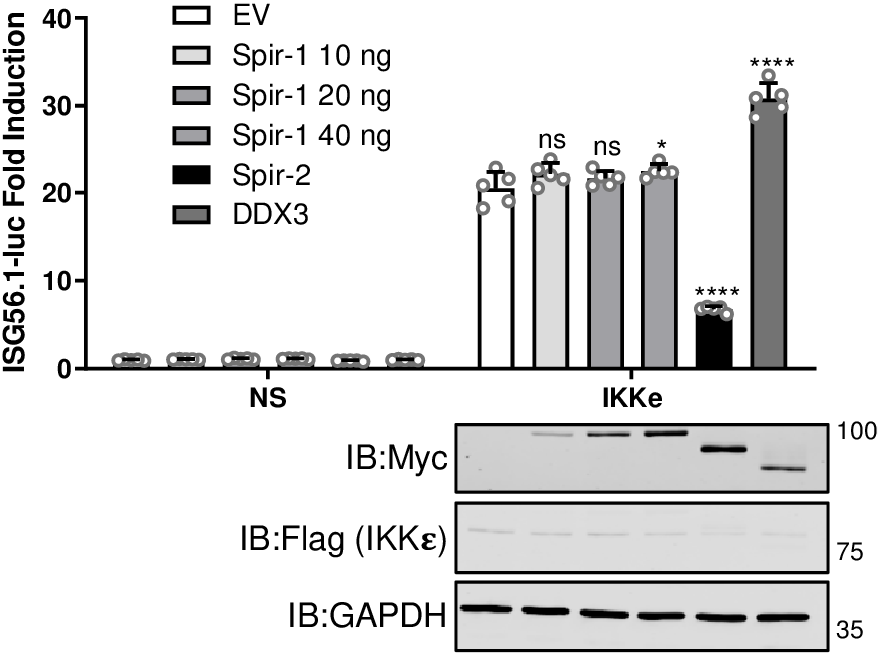
Ectopic expression of Spir-1 does not affect IRF3-dependent gene expression induced by IKKε (related to Figure 2). HEK293T cells were transfected with the ISG56.1 firefly luciferase reporter plasmid, TK-renilla luciferase and plasmids for expression of the indicated proteins. Cells were also co-transfected with EV as the non-stimulated (NS) controls or with 100 ng of IKKε plasmid to activate the IRF3 pathway. EV was added to samples when necessary to keep the final amount of DNA transfected as 40 ng in all samples. Cell lysates were prepared and luciferase expression was measured and normalised to renilla luciferase. At least triplicate samples were analysed for each condition. Data are expressed as the mean (± SEM) fold induction of the firefly luciferase activity normalised to renilla values for the stimulated versus non-stimulated samples. Data shown are representative of three independent experiments. Immunoblots underneath each graph show the expression levels of the different proteins. The positions of molecular mass markers in kDa are shown on the right and the antibodies used are shown on the left. ns = not significant; *P < 0.05; ****P < 0.0001.

**S1 Table.** Plasmids used in the study.

| PLASMID | SOURCE |
| --- | --- |
| human nMyc-Spir-1-pcDNA3.1 | Gift from Prof. Dr. E. Kerkhoff (University Hospital Regensburg, Germany) |
| human nMyc-Spir-2-pcDNA3.1 | Gift from Prof. Dr. E. Kerkhoff (University Hospital Regensburg, Germany) |
| mouse nMyc-B-TrCP-pcDNA3.1 | (Mansur *et al*, 2013) |
| GFP-Myc-pCMV | (Neidel *et al*, 2019) |
| nFlag-coK7-pcDNA4/TO | (Torres *et al*, 2016) |
| coA49-cFlag-pcDNA3.1 | (Neidel *et al.*, 2019) |
| GFP-cFlag-pcDNA4/TO | Sarah Neidel (Prof. G.L. Smith laboratory) |
| human nMyc-Spir-1-NT-pcDNA3.1 | This study |
| human nMyc-Spir-1-CT-pcDNA3.1 | This study |
| coN1-cTAP-pcDNA4/TO | (Maluquer de Motes *et al*, 2011) |
| coC6-cTAP-pcDNA4/TO | (Unterholzner *et al*, 2011) |
| nTAP-coN2-pcDNA4/TO | (Ferguson *et al*, 2013) |
| nFlag-coB14-pcDNA4/TO | (Torres *et al.*, 2016) |
| pcDNA4/TO | Invitrogen |
| nMyc-DDX3-pCMV | Gift from Dr M. Schröder - (Gu *et al*, 2013) |
| nHA-DDX3-pCMV | Gift from Dr M. Schröder - (Gu *et al.*, 2013) |
| IFN-β-luc | Gift from Dr T. Taniguchi (University of Tokyo, Japan) |
| ISRE-luc | Gift from Dr. A. Bowie (Trinity College, Dublin, Ireland) |
| NF-κB-luc | Gift from Dr. A. Bowie (Trinity College, Dublin, Ireland) |
| ISG56.1-luc | Gift from G. Sen (Cleveland Clinic, Cleveland, OH, USA). |
| TK-Renilla | Promega |
| Flag-RIG-I-CARD | Gift from A. Garcia-Sastre (Mount Sinai, New York, USA) |
| HA-MAVS | (Unterholzner *et al.*, 2011) |
| Flag-TBK1 | (Unterholzner *et al.*, 2011) |
| Flag-IKKe | (Unterholzner *et al.*, 2011) |
| IRF3-5D | (Unterholzner *et al.*, 2011) |
| human nMyc-Spir-1-FFAA-pcDNA3.1 | This study |
| pF3A-nFlag-coK7 | This study |
| pF3A-nMyc-Spir-1 | This study |
| pF3A-nMyc-DDX3 | This study |
| pF3A-F12-cFlagd | GLS - unpublished |
| pF3A-nMyc-TAB2 | GLS - unpublished |
| nFlag-coK7-D28A-pcDNA4/TO | This study |
| nFlag-coK7-D31A-pcDNA4/TO | This study |
| nFlag-coK7-DDAA-pcDNA4/TO | This study |
| pSpCas9(BB)-2A-Puro (px459) | (Ran *et al*, 2013) - Addgene plasmid #62988 |
| px459-Spir-1-sgRNA#2 | This study |
| pLKO.DCMV.TetO.mcs | (Everett *et al*, 2013) |
| pLKO.DCMV.TetO.Myc-Spir-1 | This study |
| pCMV.dR8.91 | Gift from Dr H. Laman (Lomonosov *et al*, 2011) |
| pMD-G | Gift from Dr H. Laman (Lomonosov *et al.*, 2011) |

**S2 Table.**
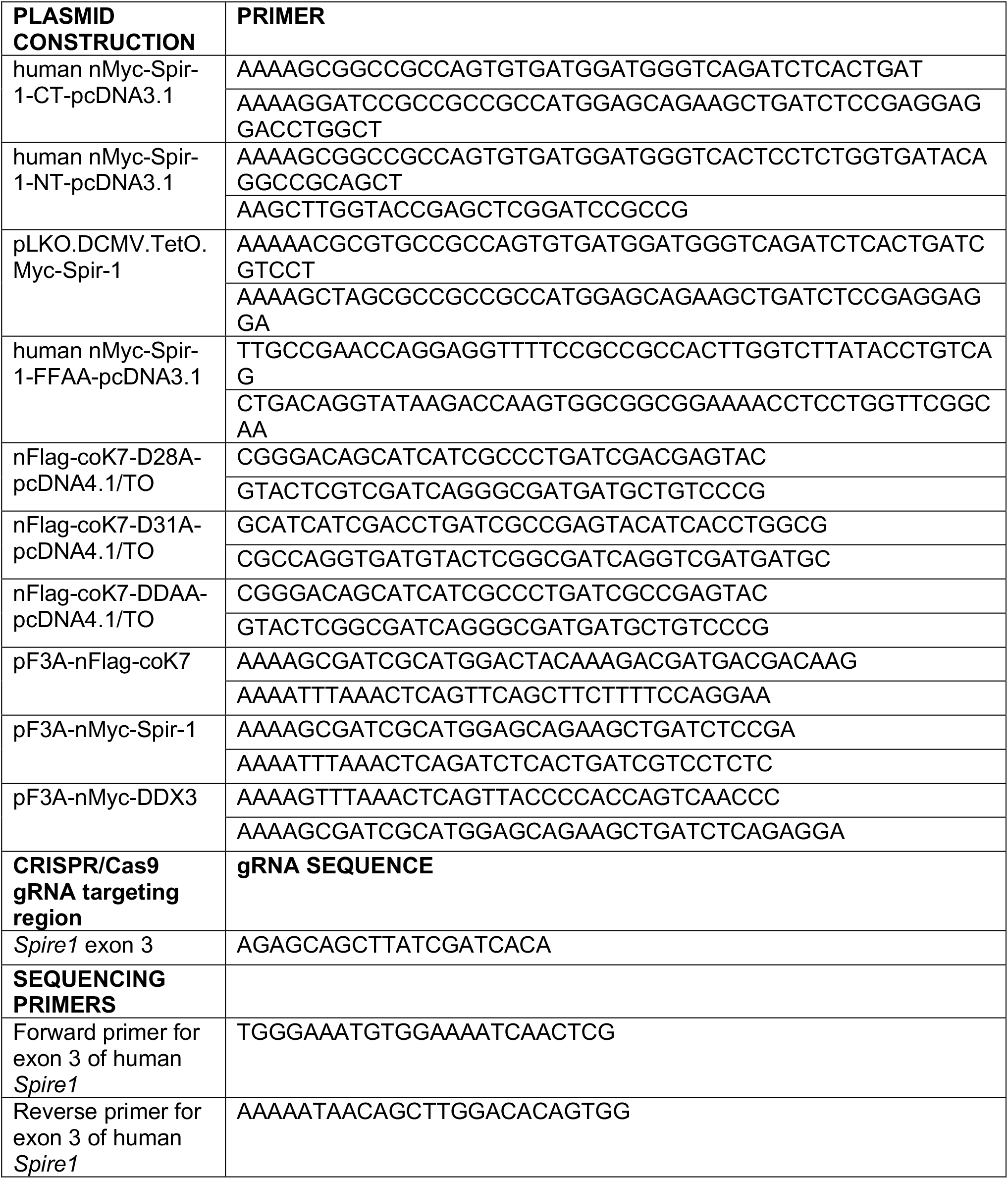
Oligonucleotides used in this study.

**S3 Table.**
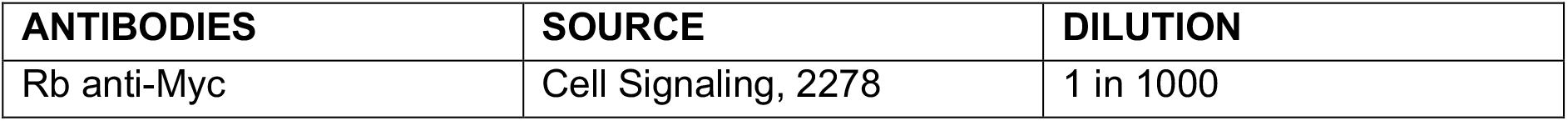
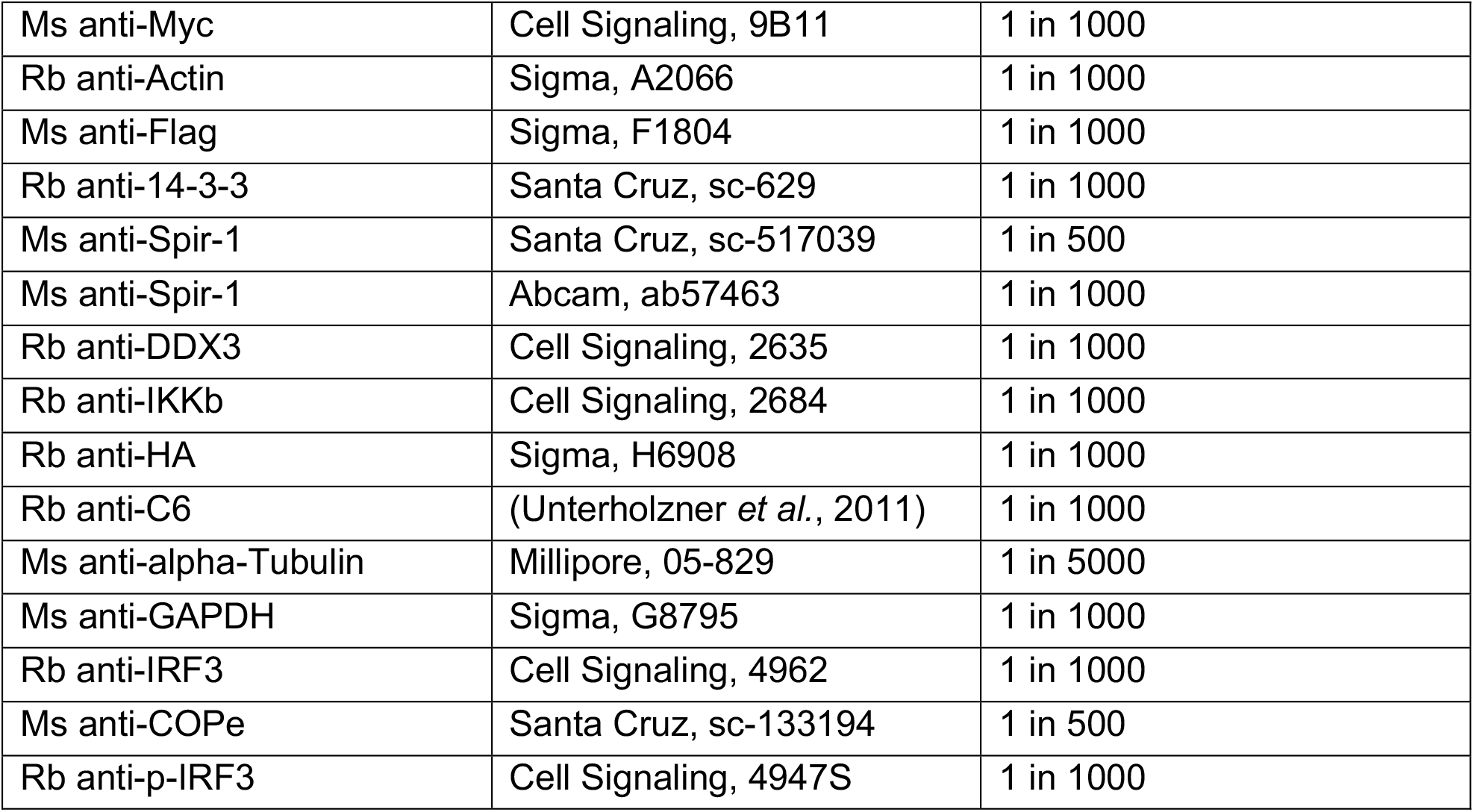
Primary antibodies used in the study.

## References

1. Janeway CA, Jr., Medzhitov R. Innate immune recognition. Annu Rev Immunol. 2002;20:197–216.

2. Pang IK, Iwasaki A. Control of antiviral immunity by pattern recognition and the microbiome. Immunol Rev. 2012;245(1):209–26.

3. Thompson MR, Kaminski JJ, Kurt-Jones EA, Fitzgerald KA. Pattern recognition receptors and the innate immune response to viral infection. Viruses. 2011;3(6):920–40.

4. Randall RE, Goodbourn S. Interferons and viruses: an interplay between induction, signalling, antiviral responses and virus countermeasures. J Gen Virol. 2008;89(Pt 1):1–47.

5. Garcia-Sastre A. Ten Strategies of Interferon Evasion by Viruses. Cell Host Microbe. 2017;22(2):176–84.

6. Fenner F, Henderson DA, Arita I, Jezek Z, Ladnyi ID. Smallpox and its eradication. World Health Organisation, Geneva. 1988.

7. Goebel SJ, Johnson GP, Perkus ME, Davis SW, Winslow JP, Paoletti E. The complete DNA sequence of vaccinia virus. Virology. 1990;179(1):247-66, 517-63.

8. Moss B, Smith GL. Poxviridae: the viruses and their replication. In: Howley PM, Knipe DM, editors. Fields Virology: DNA viruses. 2. 7th ed: Wolters Kluwer Inc; 2021. p. 573–613.

9. Smith GL, Benfield CTO, Maluquer de Motes C, Mazzon M, Ember SWJ, Ferguson BJ, et al. Vaccinia virus immune evasion: mechanisms, virulence and immunogenicity. J Gen Virol. 2013;94(Pt 11):2367–92.

10. Benfield CTO, Ren H, Lucas SJ, Bahsoun B, Smith GL. Vaccinia virus protein K7 is a virulence factor that alters the acute immune response to infection. J Gen Virol. 2013;94(Pt 7):1647–57.

11. Schroder M, Baran M, Bowie AG. Viral targeting of DEAD box protein 3 reveals its role in TBK1/IKKepsilon-mediated IRF activation. EMBO J. 2008;27(15):2147–57.

12. Teferi WM, Desaulniers MA, Noyce RS, Shenouda M, Umer B, Evans DH. The vaccinia virus K7 protein promotes histone methylation associated with heterochromatin formation. PLoS One. 2017;12(3):e0173056.

13. Li Y, Zhang L, Ke Y. Cellular interactome analysis of vaccinia virus K7 protein identifies three transport machineries as binding partners for K7. Virus Genes. 2017;53(6):814–22.

14. Pichlmair A, Kandasamy K, Alvisi G, Mulhern O, Sacco R, Habjan M, et al. Viral immune modulators perturb the human molecular network by common and unique strategies. Nature. 2012;487(7408):486–90.

15. Lu Y, Stuart JH, Talbot-Cooper C, Agrawal-Singh S, Huntly B, Smid AI, et al. Histone deacetylase 4 promotes type I interferon signaling, restricts DNA viruses, and is degraded via vaccinia virus protein C6. Proc Natl Acad Sci U S A. 2019;116(24):11997–2006.

16. Soday L, Lu Y, Albarnaz JD, Davies CTR, Antrobus R, Smith GL, et al. Quantitative Temporal Proteomic Analysis of Vaccinia Virus Infection Reveals Regulation of Histone Deacetylases by an Interferon Antagonist. Cell Rep. 2019;27(6):1920–33 e7.

17. Manseau LJ, Schupbach T. cappuccino and spire: two unique maternal-effect loci required for both the anteroposterior and dorsoventral patterns of the Drosophila embryo. Genes Dev. 1989;3(9):1437–52.

18. Wellington A, Emmons S, James B, Calley J, Grover M, Tolias P, et al. Spire contains actin binding domains and is related to ascidian posterior end mark-5. Development. 1999;126(23):5267–74.

19. Quinlan ME, Heuser JE, Kerkhoff E, Mullins RD. Drosophila Spire is an actin nucleation factor. Nature. 2005;433(7024):382–8.

20. Pechlivanis M, Samol A, Kerkhoff E. Identification of a short Spir interaction sequence at the C-terminal end of formin subgroup proteins. J Biol Chem. 2009;284(37):25324–33.

21. Bosch M, Le KH, Bugyi B, Correia JJ, Renault L, Carlier MF. Analysis of the function of Spire in actin assembly and its synergy with formin and profilin. Mol Cell. 2007;28(4):555–68.

22. Quinlan ME, Hilgert S, Bedrossian A, Mullins RD, Kerkhoff E. Regulatory interactions between two actin nucleators, Spire and Cappuccino. J Cell Biol. 2007;179(1):117–28.

23. Pfender S, Kuznetsov V, Pleiser S, Kerkhoff E, Schuh M. Spire-type actin nucleators cooperate with Formin-2 to drive asymmetric oocyte division. Curr Biol. 2011;21(11):955–60.

24. Schuh M. An actin-dependent mechanism for long-range vesicle transport. Nat Cell Biol. 2011;13(12):1431–6.

25. Vizcarra CL, Kreutz B, Rodal AA, Toms AV, Lu J, Zheng W, et al. Structure and function of the interacting domains of Spire and Fmn-family formins. Proc Natl Acad Sci U S A. 2011;108(29):11884–9.

26. Zeth K, Pechlivanis M, Samol A, Pleiser S, Vonrhein C, Kerkhoff E. Molecular basis of actin nucleation factor cooperativity: crystal structure of the Spir-1 kinase non-catalytic C-lobe domain (KIND)*formin-2 formin SPIR interaction motif (FSI) complex. J Biol Chem. 2011;286(35):30732–9.

27. Quinlan ME. Direct interaction between two actin nucleators is required in Drosophila oogenesis. Development. 2013;140(21):4417–25.

28. Montaville P, Jegou A, Pernier J, Compper C, Guichard B, Mogessie B, et al. Spire and Formin 2 synergize and antagonize in regulating actin assembly in meiosis by a ping-pong mechanism. PLoS Biol. 2014;12(2):e1001795.

29. Tittel J, Welz T, Czogalla A, Dietrich S, Samol-Wolf A, Schulte M, et al. Membrane targeting of the Spir.formin actin nucleator complex requires a sequential handshake of polar interactions. J Biol Chem. 2015;290(10):6428–44.

30. Ducka AM, Joel P, Popowicz GM, Trybus KM, Schleicher M, Noegel AA, et al. Structures of actin-bound Wiskott-Aldrich syndrome protein homology 2 (WH2) domains of Spire and the implication for filament nucleation. Proc Natl Acad Sci U S A. 2010;107(26):11757–62.

31. Sitar T, Gallinger J, Ducka AM, Ikonen TP, Wohlhoefler M, Schmoller KM, et al. Molecular architecture of the Spire-actin nucleus and its implication for actin filament assembly. Proc Natl Acad Sci U S A. 2011;108(49):19575–80.

32. Pylypenko O, Welz T, Tittel J, Kollmar M, Chardon F, Malherbe G, et al. Coordinated recruitment of Spir actin nucleators and myosin V motors to Rab11 vesicle membranes. Elife. 2016;5.

33. Alzahofi N, Welz T, Robinson CL, Page EL, Briggs DA, Stainthorp AK, et al. Rab27a coordinates actin-dependent transport by controlling organelle-associated motors and track assembly proteins. Nat Commun. 2020;11(1):3495.

34. Kerkhoff E, Simpson JC, Leberfinger CB, Otto IM, Doerks T, Bork P, et al. The Spir actin organizers are involved in vesicle transport processes. Curr Biol. 2001;11(24):1963–8.

35. Lagal V, Abrivard M, Gonzalez V, Perazzi A, Popli S, Verzeroli E, et al. Spire-1 contributes to the invadosome and its associated invasive properties. J Cell Sci. 2014;127(Pt 2):328–40.

36. Schumacher N, Borawski JM, Leberfinger CB, Gessler M, Kerkhoff E. Overlapping expression pattern of the actin organizers Spir-1 and formin-2 in the developing mouse nervous system and the adult brain. Gene Expr Patterns. 2004;4(3):249–55.

37. Pleiser S, Rock R, Wellmann J, Gessler M, Kerkhoff E. Expression patterns of the mouse Spir-2 actin nucleator. Gene Expr Patterns. 2010;10(7-8):345–50.

38. Fagerberg L, Hallstrom BM, Oksvold P, Kampf C, Djureinovic D, Odeberg J, et al. Analysis of the human tissue-specific expression by genome-wide integration of transcriptomics and antibody-based proteomics. Mol Cell Proteomics. 2014;13(2):397–406.

39. Morel E, Parton RG, Gruenberg J. Annexin A2-dependent polymerization of actin mediates endosome biogenesis. Dev Cell. 2009;16(3):445–57.

40. Belin BJ, Lee T, Mullins RD. DNA damage induces nuclear actin filament assembly by Formin - 2 and Spire-(1/2) that promotes efficient DNA repair. [corrected]. Elife. 2015;4:e07735.

41. Manor U, Bartholomew S, Golani G, Christenson E, Kozlov M, Higgs H, et al. A mitochondria-anchored isoform of the actin-nucleating spire protein regulates mitochondrial division. Elife. 2015;4.

42. Wen Q, Li N, Xiao X, Lui WY, Chu DS, Wong CKC, et al. Actin nucleator Spire 1 is a regulator of ectoplasmic specialization in the testis. Cell Death Dis. 2018;9(2):208.

43. Ovsyannikova IG, Kennedy RB, O’Byrne M, Jacobson RM, Pankratz VS, Poland GA. Genome-wide association study of antibody response to smallpox vaccine. Vaccine. 2012;30(28):4182–9.

44. Kerkhoff E. Cellular functions of the Spir actin-nucleation factors. Trends Cell Biol. 2006;16(9):477–83.

45. Kollmar M, Welz T, Straub F, Alzahofi N, Hatje K, Briggs DA, et al. Animal evolution coincides with a novel degree of freedom in exocytic transport processes. bioRxiv. 2020;https://doi.org/10.1101/591974.

46. Neidel S, Maluquer de Motes C, Mansur DS, Strnadova P, Smith GL, Graham SC. Vaccinia virus protein A49 is an unexpected member of the B-cell Lymphoma (Bcl)-2 protein family. J Biol Chem. 2015;290(10):5991–6002.

47. Mansur DS, Maluquer de Motes C, Unterholzner L, Sumner RP, Ferguson BJ, Ren H, et al. Poxvirus targeting of E3 ligase beta-TrCP by molecular mimicry: a mechanism to inhibit NF-kappaB activation and promote immune evasion and virulence. PLoS Pathog. 2013;9(2):e1003183.

48. Benzinger A, Muster N, Koch HB, Yates JR, 3rd, Hermeking H. Targeted proteomic analysis of 14-3-3 sigma, a p53 effector commonly silenced in cancer. Mol Cell Proteomics. 2005;4(6):785–95.

49. Ewing RM, Chu P, Elisma F, Li H, Taylor P, Climie S, et al. Large-scale mapping of human protein-protein interactions by mass spectrometry. Mol Syst Biol. 2007;3:89.

50. Chen RA, Jacobs N, Smith GL. Vaccinia virus strain Western Reserve protein B14 is an intracellular virulence factor. J Gen Virol. 2006;87(Pt 6):1451–8.

51. Chen RA, Ryzhakov G, Cooray S, Randow F, Smith GL. Inhibition of IkappaB kinase by vaccinia virus virulence factor B14. PLoS Pathog. 2008;4(2):e22.

52. Kato H, Takeuchi O, Sato S, Yoneyama M, Yamamoto M, Matsui K, et al. Differential roles of MDA5 and RIG-I helicases in the recognition of RNA viruses. Nature. 2006;441(7089):101–5.

53. Unterholzner L, Sumner RP, Baran M, Ren H, Mansur DS, Bourke NM, et al. Vaccinia virus protein C6 is a virulence factor that binds TBK-1 adaptor proteins and inhibits activation of IRF3 and IRF7. PLoS Pathog. 2011;7(9):e1002247.

54. Stuart JH, Sumner RP, Lu Y, Snowden JS, Smith GL. Vaccinia Virus Protein C6 Inhibits Type I IFN Signalling in the Nucleus and Binds to the Transactivation Domain of STAT2. PLoS Pathog. 2016;12(12):e1005955.

55. Ferguson BJ, Benfield CTO, Ren H, Lee VH, Frazer GL, Strnadova P, et al. Vaccinia virus protein N2 is a nuclear IRF3 inhibitor that promotes virulence. J Gen Virol. 2013;94(Pt 9):2070–81.

56. Kalverda AP, Thompson GS, Vogel A, Schroder M, Bowie AG, Khan AR, et al. Poxvirus K7 protein adopts a Bcl-2 fold: biochemical mapping of its interactions with human DEAD box RNA helicase DDX3. J Mol Biol. 2009;385(3):843–53.

57. Oda S, Schroder M, Khan AR. Structural basis for targeting of human RNA helicase DDX3 by poxvirus protein K7. Structure. 2009;17(11):1528–37.

58. Fullam A, Schroder M. DExD/H-box RNA helicases as mediators of anti-viral innate immunity and essential host factors for viral replication. Biochim Biophys Acta. 2013;1829(8):854–65.

59. Gu L, Fullam A, McCormack N, Hohn Y, Schroder M. DDX3 directly regulates TRAF3 ubiquitination and acts as a scaffold to co-ordinate assembly of signalling complexes downstream from MAVS. Biochem J. 2017;474(4):571–87.

60. Wang T, Birsoy K, Hughes NW, Krupczak KM, Post Y, Wei JJ, et al. Identification and characterization of essential genes in the human genome. Science. 2015;350(6264):1096–101.

61. Carpentier DC, Gao WN, Ewles H, Morgan GW, Smith GL. Vaccinia virus protein complex F12/E2 interacts with kinesin light chain isoform 2 to engage the kinesin-1 motor complex. PLoS Pathog. 2015;11(3):e1004723.

62. Everett RD, Parsy ML, Orr A. Analysis of the functions of herpes simplex virus type 1 regulatory protein ICP0 that are critical for lytic infection and derepression of quiescent viral genomes. J Virol. 2009;83(10):4963–77.

63. Stinski MF, Meier JL. Immediate-early viral gene regulation and function. In: Arvin A, Campadelli-Fiume G, Mocarski E, Moore PS, Roizman B, Whitley R, et al., editors. Human Herpesviruses: Biology, Therapy, and Immunoprophylaxis. Cambridge 2007.

64. Servant MJ, Grandvaux N, tenOever BR, Duguay D, Lin R, Hiscott J. Identification of the minimal phosphoacceptor site required for in vivo activation of interferon regulatory factor 3 in response to virus and double-stranded RNA. J Biol Chem. 2003;278(11):9441–7.

65. Pleiser S, Banchaabouchi MA, Samol-Wolf A, Farley D, Welz T, Wellbourne-Wood J, et al. Enhanced fear expression in Spir-1 actin organizer mutant mice. Eur J Cell Biol. 2014;93(5-6):225–37.

66. Carter GC, Rodger G, Murphy BJ, Law M, Krauss O, Hollinshead M, et al. Vaccinia virus cores are transported on microtubules. J Gen Virol. 2003;84(Pt 9):2443–58.

67. Neidel S, Torres AA, Ren H, Smith GL. Leaky scanning translation generates a second A49 protein that contributes to vaccinia virus virulence. J Gen Virol. 2020;101(5):533–41.

68. Smith GL, Vanderplasschen A, Law M. The formation and function of extracellular enveloped vaccinia virus. J Gen Virol. 2002;83(Pt 12):2915–31.

69. Kato H, Sato S, Yoneyama M, Yamamoto M, Uematsu S, Matsui K, et al. Cell type-specific involvement of RIG-I in antiviral response. Immunity. 2005;23(1):19–28.

70. Kato H, Takeuchi O, Mikamo-Satoh E, Hirai R, Kawai T, Matsushita K, et al. Length-dependent recognition of double-stranded ribonucleic acids by retinoic acid-inducible gene-I and melanoma differentiation-associated gene 5. J Exp Med. 2008;205(7):1601–10.

71. Brisse M, Ly H. Comparative Structure and Function Analysis of the RIG-I-Like Receptors: RIG-I and MDA5. Front Immunol. 2019;10:1586.

72. Soulat D, Burckstummer T, Westermayer S, Goncalves A, Bauch A, Stefanovic A, et al. The DEAD-box helicase DDX3X is a critical component of the TANK-binding kinase 1-dependent innate immune response. EMBO J. 2008;27(15):2135–46.

73. Seth RB, Sun L, Ea CK, Chen ZJ. Identification and characterization of MAVS, a mitochondrial antiviral signaling protein that activates NF-kappaB and IRF 3. Cell. 2005;122(5):669–82.

74. Weber F, Wagner V, Rasmussen SB, Hartmann R, Paludan SR. Double-stranded RNA is produced by positive-strand RNA viruses and DNA viruses but not in detectable amounts by negative-strand RNA viruses. J Virol. 2006;80(10):5059–64.

75. Liu Y, Olagnier D, Lin R. Host and Viral Modulation of RIG-I-Mediated Antiviral Immunity. Front Immunol. 2016;7:662.

76. Serman TM, Gack MU. Evasion of Innate and Intrinsic Antiviral Pathways by the Zika Virus. Viruses. 2019;11(10).

77. Colby C, Duesberg PH. Double-stranded RNA in vaccinia virus infected cells. Nature. 1969;222(5197):940–4.

78. Rice AP, Roberts WK, Kerr IM. 2-5A accumulates to high levels in interferon-treated, vaccinia virus-infected cells in the absence of any inhibition of virus replication. J Virol. 1984;50(1):220–8.

79. Pichlmair A, Schulz O, Tan CP, Rehwinkel J, Kato H, Takeuchi O, et al. Activation of MDA5 requires higher-order RNA structures generated during virus infection. J Virol. 2009;83(20):10761–9.

80. Marq JB, Hausmann S, Luban J, Kolakofsky D, Garcin D. The double-stranded RNA binding domain of the vaccinia virus E3L protein inhibits both RNA- and DNA-induced activation of interferon beta. J Biol Chem. 2009;284(38):25471–8.

81. Valentine R, Smith GL. Inhibition of the RNA polymerase III-mediated dsDNA-sensing pathway of innate immunity by vaccinia virus protein E3. J Gen Virol. 2010;91(Pt 9):2221–9.

82. Liu SW, Katsafanas GC, Liu R, Wyatt LS, Moss B. Poxvirus decapping enzymes enhance virulence by preventing the accumulation of dsRNA and the induction of innate antiviral responses. Cell Host Microbe. 2015;17(3):320–31.

83. Mutso M, Saul S, Rausalu K, Susova O, Zusinaite E, Mahalingam S, et al. Reverse genetic system, genetically stable reporter viruses and packaged subgenomic replicon based on a Brazilian Zika virus isolate. J Gen Virol. 2017;98(11):2712–24.

84. Torres AA, Albarnaz JD, Bonjardim CA, Smith GL. Multiple Bcl-2 family immunomodulators from vaccinia virus regulate MAPK/AP-1 activation. J Gen Virol. 2016;97(9):2346–51.

85. Odon V, Georgana I, Holley J, Morata J, Maluquer de Motes C. Novel Class of Viral Ankyrin Proteins Targeting the Host E3 Ubiquitin Ligase Cullin-2. J Virol. 2018;92(23).

86. Everett RD, Bell AJ, Lu Y, Orr A. The replication defect of ICP0-null mutant herpes simplex virus 1 can be largely complemented by the combined activities of human cytomegalovirus proteins IE1 and pp71. J Virol. 2013;87(2):978–90.

87. Ran FA, Hsu PD, Wright J, Agarwala V, Scott DA, Zhang F. Genome engineering using the CRISPR-Cas9 system. Nat Protoc. 2013;8(11):2281–308.

## References

Everett RD, Bell AJ, Lu Y, Orr A (2013) The replication defect of ICP0-null mutant herpes simplex virus 1 can be largely complemented by the combined activities of human cytomegalovirus proteins IE1 and pp71. J Virol 87: 978–990

Ferguson BJ, Benfield CTO, Ren H, Lee VH, Frazer GL, Strnadova P, Sumner RP, Smith GL (2013) Vaccinia virus protein N2 is a nuclear IRF3 inhibitor that promotes virulence. J Gen Virol 94: 2070–2081

Gu L, Fullam A, Brennan R, Schroder M (2013) Human DEAD box helicase 3 couples IkappaB kinase epsilon to interferon regulatory factor 3 activation. Mol Cell Biol 33: 2004–2015

Lomonosov M, Meziane el K, Ye H, Nelson DE, Randle SJ, Laman H (2011) Expression of Fbxo7 in haematopoietic progenitor cells cooperates with p53 loss to promote lymphomagenesis. PLoS One 6: e21165

Maluquer de Motes C, Cooray S, Ren H, Almeida GM, McGourty K, Bahar MW, Stuart DI, Grimes JM, Graham SC, Smith GL (2011) Inhibition of apoptosis and NF-kappaB activation by vaccinia protein N1 occur via distinct binding surfaces and make different contributions to virulence. PLoS Pathog 7: e1002430

Mansur DS, Maluquer de Motes C, Unterholzner L, Sumner RP, Ferguson BJ, Ren H, Strnadova P, Bowie AG, Smith GL (2013) Poxvirus targeting of E3 ligase beta-TrCP by molecular mimicry: a mechanism to inhibit NF-kappaB activation and promote immune evasion and virulence. PLoS Pathog 9: e1003183

Neidel S, Ren H, Torres AA, Smith GL (2019) NF-kappaB activation is a turn on for vaccinia virus phosphoprotein A49 to turn off NF-kappaB activation. Proc Natl Acad Sci U S A 116: 5699–5704

Ran FA, Hsu PD, Wright J, Agarwala V, Scott DA, Zhang F (2013) Genome engineering using the CRISPR-Cas9 system. Nat Protoc 8: 2281–2308

Torres AA, Albarnaz JD, Bonjardim CA, Smith GL (2016) Multiple Bcl-2 family immunomodulators from vaccinia virus regulate MAPK/AP-1 activation. J Gen Virol 97: 2346–2351

Unterholzner L, Sumner RP, Baran M, Ren H, Mansur DS, Bourke NM, Randow F, Smith GL, Bowie AG (2011) Vaccinia virus protein C6 is a virulence factor that binds TBK-1 adaptor proteins and inhibits activation of IRF3 and IRF7. PLoS Pathog 7: e1002247

